# A comparative study of deconvolution methods for RNA-seq data under a dynamic testing landscape

**DOI:** 10.1101/2020.12.09.418640

**Authors:** Haijing Jin, Zhandong Liu

## Abstract

Deconvolution analyses have been widely used to track compositional alternations of cell-types in gene expression data. Even though numerous novel methods have been developed in recent years, researchers are still having difficulty selecting optimal deconvolution methods due to the lack of comprehensive benchmarks relative to the newly developed methods. To systematically reveal the pitfalls and challenges of deconvolution analyses, we studied the impact of several technical and biological factors such as simulation model, quantification unit, component number, weight matrix, and unknown content by constructing three benchmarking frameworks that cover comparative analysis of 11 popular deconvolution methods under 1,766 conditions. We hope this study can provide new insights to researchers for future application, standardization, and development of deconvolution tools on RNA-seq data.

## Background

Deconvolution refers to a process that separates a heterogeneous mixture signal into its constituent components. In the biomedical field, researchers have been using deconvolution methods to derive cell-type-specific signals^1–3^ from heterogeneous mixture data. Cellular composition information is crucial for developing sophisticated diagnostic techniques as it enables researchers to track each cellular component’s contribution during disease progressions^4^. Although some experimental approaches like fluorescence-activated cell sorting(FACS), immunohistochemistry(IHC), and single-cell RNA-seq can derive cell-type proportion data^3^, all these approaches are either restricted by its throughput or remain too costly and laborious for large-scale clinical applications. By far, deconvolution is recognized as the most cost-effect approach to derive cell-type proportion data from heterogenous biospecimens and has the potential to bring a considerable improvement in the speed and scale of cell-type-specific clinical diagnosis.

By January 2018, there have been around 50 deconvolution methods developed^2^ and researchers are now facing the challenge of selecting the right method for deconvolution analysis. In a methodological paper, authors usually compared the method of their own to a chosen set of published methods and arrived at the conclusion that their method was the best. However, only a limited number of deconvolution methods and biological conditions were considered in these comparisons. Moreover, different research groups applied inconsistent testing frameworks with different simulation strategies, evaluation metrics, and cell-type annotations, making it difficult for researchers to determine the optimal method for the deconvolution analysis. For a fair and comprehensive comparison of deconvolution applications in complex biological systems, an independent benchmarking is in need^5^. Previously, Sturm *et al*.^3^ and Cobos *et al*.^6^ performed quantitative evaluations of reference-based and marker-based deconvolution methods on RNA-seq data. Sturm *et al*.^3^ focused on spill-over effects, minimal detection fraction, and background predictions and suggested removing non-specific signature genes to improve deconvolution accuracy. Cobos *et al*.^6^ focused on the impact of different normalization strategies, reference platforms, marker gene selection strategies, and missing cellular components in the reference. Compared with previous benchmarks, our study focuses on technical and biological factors caused by varied experimental mixture conditions such as mixture noise levels, quantification unit selection, cellular component number, weight matrix property, and unknown cellular contents. We also studied the major factors that determine an evaluation framework, such as simulation model selection, evaluation metric selection, and measurement scale selection. Our work carefully examined the joint impact of different technical parameters and biological design factors to provide an insightful reference guide for mixture condition determination and deconvolution method selection.

There are three types of benchmarking frameworks for the evaluation of deconvolution methods: *in silico* framework^7,8^, *in vitro* framework^9^, and *in vivo* framework^10^ (Supplementary Table 1). The *in vivo* testing framework mainly rely on indirect performance assessment and usually cannot derive a definite conclusion of the method’s performance. Only a few in vivo benchmarking datasets^3^ have coupled FACS results. Nevertheless, these benchmarking datasets are often restricted by limited cell types and sample numbers^3,8^. The *in vitro* testing framework where mixtures are generated in the tube with predefined mixing compositions also suffers from limited cell types and sample numbers. Moreover, most *in vitro* testing frameworks applied ‘orthogonal’ weights, leading to over-optimistic performance assessment. The *in silico* testing framework uses RNA-seq profiles from purified biological samples as primary building blocks and generates heterogeneous mixing samples by *in silico* mixing procedures. Among all three benchmarking frameworks, we selected the *in silico* testing framework to systematically explore the impact of different biological and technical factors, which require large amounts of benchmarking datasets under controlled and finely tuned multi-factor testing environments.

To provide a reliable reference for the application and development of deconvolution methods, we compared 11 deconvolution methods (Figure 1b and Supplementary Table 3). To establish benchmarking frameworks that mimic application scenarios of more complicated and diverse biological systems, we designed three sets of benchmarking frameworks that mimic up to 1,766 biological conditions with varying noise levels, library sizes, cellular component numbers, weight matrix properties, simulation models, and proportions of unknown contents (Figure 1a, Supplementary Table 2). To determine the impact of evaluation frameworks, we performed comparisons under different simulation models and measurement scales with two sets of evaluation metrics: correlation (Pearson’s Correlation Coefficient) and mAD (Mean Absolute Deviation)(Methods). Compared with previous benchmarks, we applied more flexible and sophisticated simulation strategies to create mixtures covering dynamic conditions, which enable us to investigate the tipping point where each method deteriorates. Moreover, we studied the impact of commonly applied simulation strategies, and by comparison to the real mixture data, we derived improved simulation strategies that can generate more complex and yet authentic simulation data. Our results provide a dynamic testing landscape that allows the user to select the right method that performs well in the targeted experimental condition.

**Fig.1.**
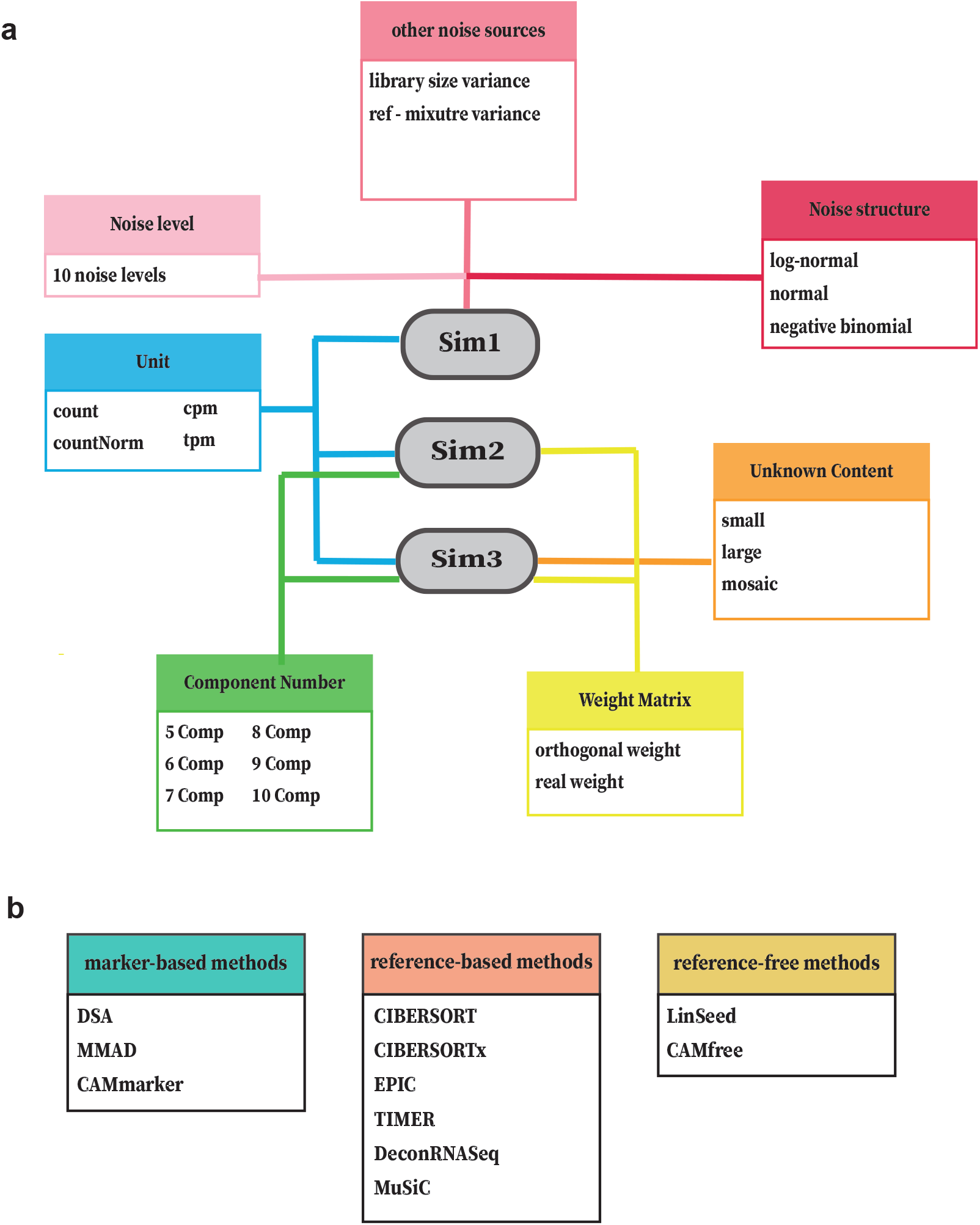
Overview of *in silico* testing frameworks and methods categorization. **a**, Three benchmarking frameworks were constructed to investigate the impact of seven factors that affect deconvolution analysis: noise level, noise structure, other noise sources, quantification unit, unknown content, component number, and weight matrix. **b**, 11 deconvolution methods are tested and have been categorized based on the required reference input: marker-based, reference-based, and reference-free.

## Results

### Using simulation to generate diverse deconvolution testing environments

We designed three benchmarking frameworks to test the performance of deconvolution methods under multiple application scenarios. Each framework was designed to study the impact of specific technical and biological factors on deconvolution analysis (Figure 1a). The first benchmarking framework (Sim1) was designed to reveal the impact of the noise structure under diverse noise levels. The second benchmarking framework (Sim2) was designed to reveal the impact of cellular component numbers and weight matrix properties. The third benchmarking framework (Sim3) was designed to reveal the impact of unknown biological contents and measurement scales.

In an *in silico* benchmarking framework, a deconvolution testing environment consists of mixture data, reference data, ground truths, and testing methods. Mixture data refers to heterogeneous gene expression profiles for deconvolution. Reference data refers to homogeneous cell-type-specific data, which is used to guide the deconvolution process.

Ground truths refer to the real mixing proportions of constituent cell types in the mixture data. The accuracy of deconvolution methods can be assessed by comparing estimated proportions to the ground truths. Reference data can vary based on the required input of the tested deconvolution method. In this study, we classified eleven deconvolution methods according to the required reference data in the following categories: marker-based, reference-based, and reference-free (Figure 1b, Supplementary Table 3). Marker-based methods such as DSA^11^, MMAD^12^, and CAMmarker^13^ use marker gene lists to guide the deconvolution analysis. Reference-based methods such as CIBERSORT^7^, CIBERSORTx^8^, EPIC^14^, TIMER^10^, DeconRNASeq^15^, and MuSiC^16^ use cell-type-specific gene expression profiles. Except for MuSiC^16^, nearly all reference-based methods require signature gene lists as an additional input. MuSiC^16^ implements weighted non-negative least squares regression (W-NNLS) and does not require any pre-determined gene sets. Finally, reference-free methods such as LinSeed^17^ and CAMfree^13^ do not require any external references.

### Selection of simulation model affects the deconvolution evaluation

The benchmarking framework Sim1_simModel is designed to learn the impact of noise structure under different noise levels (Fig. 1a, Methods). To understand the impact of noise structure, we simulated noise based on three simulation models: normal, log-normal, and negative binomial (nb). All these simulation models have been applied in previous publications^7,15,17–19^ to generate *in silico* mixing expression profiles. For each simulation model, we generated ten levels of noise to evaluate the robustness of deconvolution methods to the magnitude of noise(Supplementary Fig. 1a). To ensure the generality of our conclusion across different datasets and account for reference-mixture variance, we performed repeated mixture simulation with three independent blood datasets and created nine testing environments with different mixture-reference pairs (Methods, Supplementary Table 2 and Supplementary Table 4).

For the noise level, consistent with previous findings, we observed that the accuracies of the deconvolution methods decreased as the noise level increased, which was exhibited as decreasing correlation (Supplementary Fig. 3) and increasing mAD (Supplementary Fig. 4) values. We also noticed that the impact of the RNA-seq quantification unit is trivial (Supplementary Fig. 3 and 4) and thus selected the most commonly used unit tpm for remaining illustrations of testing results in Sim1_simModel. Unless specifically indicated (as in Sim1_libSize), all results in this study are from mixture data with the tpm unit.

To reveal the impact of the simulation models, we averaged evaluation metrics across noise levels and generated summarized evaluation heatmaps (11 × 3) where row index number 11 indicates the number of methods and column index number 3 indicates the number of simulation models. Based on the summarized evaluation heatmaps of correlation (Fig. 2a) and mAD (Supplementary Fig. 5a), we observed that the selection of the simulation model strongly affected evaluation results. For instance, methods like DSA^11^, TIMER^10^, and CAMfree’s^13^ rankings were all relatively higher in the negative binomial group in both correlation (Fig. 2b) and mAD (Supplementary Fig. 5b) metrics when comparing with evaluations from normal and log-normal groups. The above phenomenon indicated that the performance of some deconvolution methods is underestimated due to the underlying simulation model.

**Fig.2.**
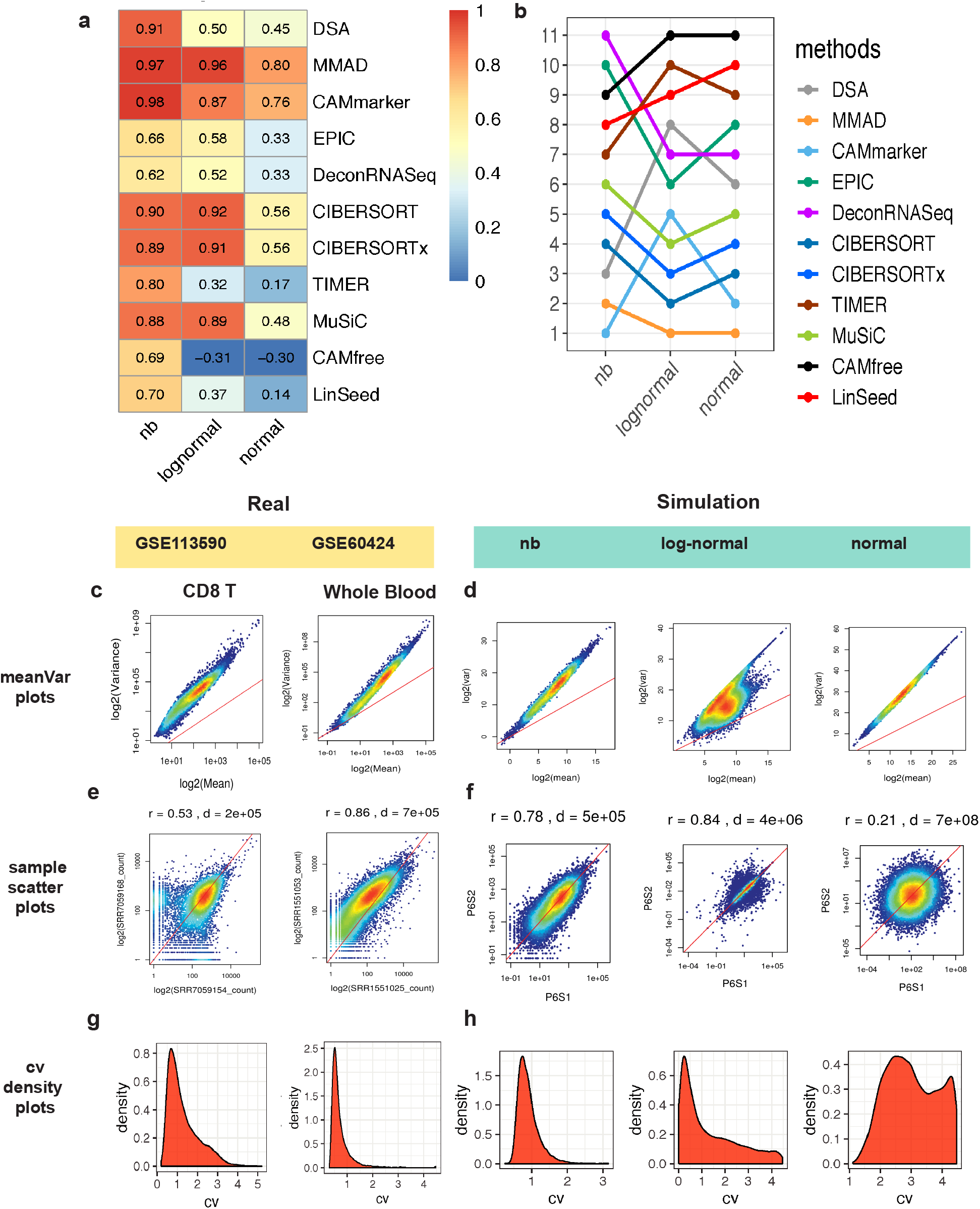
Evaluation results of Sim1_simModel and noise structure comparisons between real and simulated data. **a**, Heatmap of summarized evaluation results based on the Pearson’s correlation coefficients and **b**, rankings of tested deconvolution methods in the Sim1_simModel. In each heatmap, row indexes refer to the tested methods and column indexes refer to the simulation models (negative binomial, log-normal, and normal). **c,d**, Mean-variance plots of (**c**) real and (**d**) simulated data. (r: Spearman’s correlation coefficient, d: Euclidean distance) **e,f**, sample-sample scatter plots of (**e**) real and (**f**) simulated data. **g,h**, Density plots of CV (Coefficient of variation) of (**g**) real and (**d**) simulated data. (Real data are derived from GSE113590 and GSE60424 and Supplementary Figure 6 and 7 contain detailed variance analysis results for each dataset) (All simulated data in Figure 2 are based on simulations derived from GSE51984 with the P6 noise level.) (Results in **a** and **b** are in tpm unit, results in **c-f** are in count unit)

### The negative binomial model recapitulates noise structures of real data

In the Sim1_simModel, we found that the noise structure is the main factor obscuring deconvolution performance assessment (Fig. 2a and b, Supplementary Fig. 5). To identify the simulation model that best recapitulates the essential characteristics of real data, we performed noise structure comparisons between real and simulated data by mean-variance plots, sample-sample scatter plots and coefficient of variance (CV) density plots.

We used the mean-variance plots to study the overall trend of variance along with the gene expression level in both real and simulated data (P6 noise level) (Fig. 2c and d). As expected, we observed that the variance and mean value of counts follow a linear trend in the log space with a clear overdispersion phenomenon, which is typical to the RNA-seq data^20^(Fig. 2c). However, in the simulation group, only the simulations generated from the negative binomial and normal models showed a similar mean-variance trend to the trend observed in the real data (Fig. 2d).

Next, we used sample-sample scatter plots to study the concordance trend of gene expression profiles(Fig. 2e and f). In real data, we observed that lowly expressed genes exhibited larger relative deviances to the diagonal reference line (y = x) than highly expressed genes (Fig. 2e). This phenomenon indicates larger uncertainties in quantifying RNA molecules with lower abundance. In the simulation group, only simulation data from the negative binomial model recapitulated higher deviances of lowly expressed genes (Fig. 2f).

We also compared the magnitude of noise between the real and simulated data. In the real data, the sample-sample Spearman’s correlation values range from 0.53 to 0.99 while the sample-sample Euclidean distances fluctuate around the order of 10^4^~ 10^5^ (Supplementary Fig.6 a and b and Supplementary Fig. 7 a and b). In three tested simulation models, only the negative binomial model was capable of generating simulated profiles with comparable sample-sample correlation (0.57 - 0.98) and Euclidean distance (around the order of 10^4^~ 10^5^) to the real datasets (Supplementary Figure 8) while maintaining mean-variance trend with overdispersion phenomenon (Supplementary Fig. 9).

We compared the density curve of CV (coefficient variation) values in real and simulated data (Fig. 2g and h). Real data exhibited a unimodal bell-shaped curve, indicating that most of the genes had low to moderate levels of CV (Fig. 2g). In the simulation group, only simulations derived from the negative binomial model maintained the unimodal bell-shaped curve throughout all noise levels (Fig. 2h). CV density distributions of normal and log-normal simulation models showed density curves that were skewed towards the high CV value from noise level P6 to P10, which indicating unauthentic noise structure(Supplementary Fig. 10b).

In conclusion, the negative binomial simulation model, which successfully recapitulates the mean-variance trend, sample-sample concordance, the density of CV, presents the most similar noise structure to the real data. The negative binomial model also kept the magnitude of noise at comparable levels to the real data and thus should be considered as the most appropriate simulation model for generating *in silico* mixtures for deconvolution benchmarking.

### Library size normalization is required to ensure the deconvolution accuracy

In this benchmarking framework, we focused on the impact of RNA-seq quantification units with mixtures that varied in their library sizes (Supplementary Fig. 1b). To reveal bias caused by varied library sizes, we designed Sim1_libSize in which every mixture comprised of samples with varied library sizes (first 10 samples with 12M reads, and remaining 10 samples with 24M reads), and our results indicate using quantification units normalized by library sizes can mitigate the bias caused by library size variation (Fig. 3a, Supplementary Fig. 11a). We summarized evaluation results across all 10 noise levels and generated evaluation heatmaps with dimensions 11 by 4 where 11 indicates the number of methods and 4 indicates the number of quantification units being tested.

**Fig.3.**
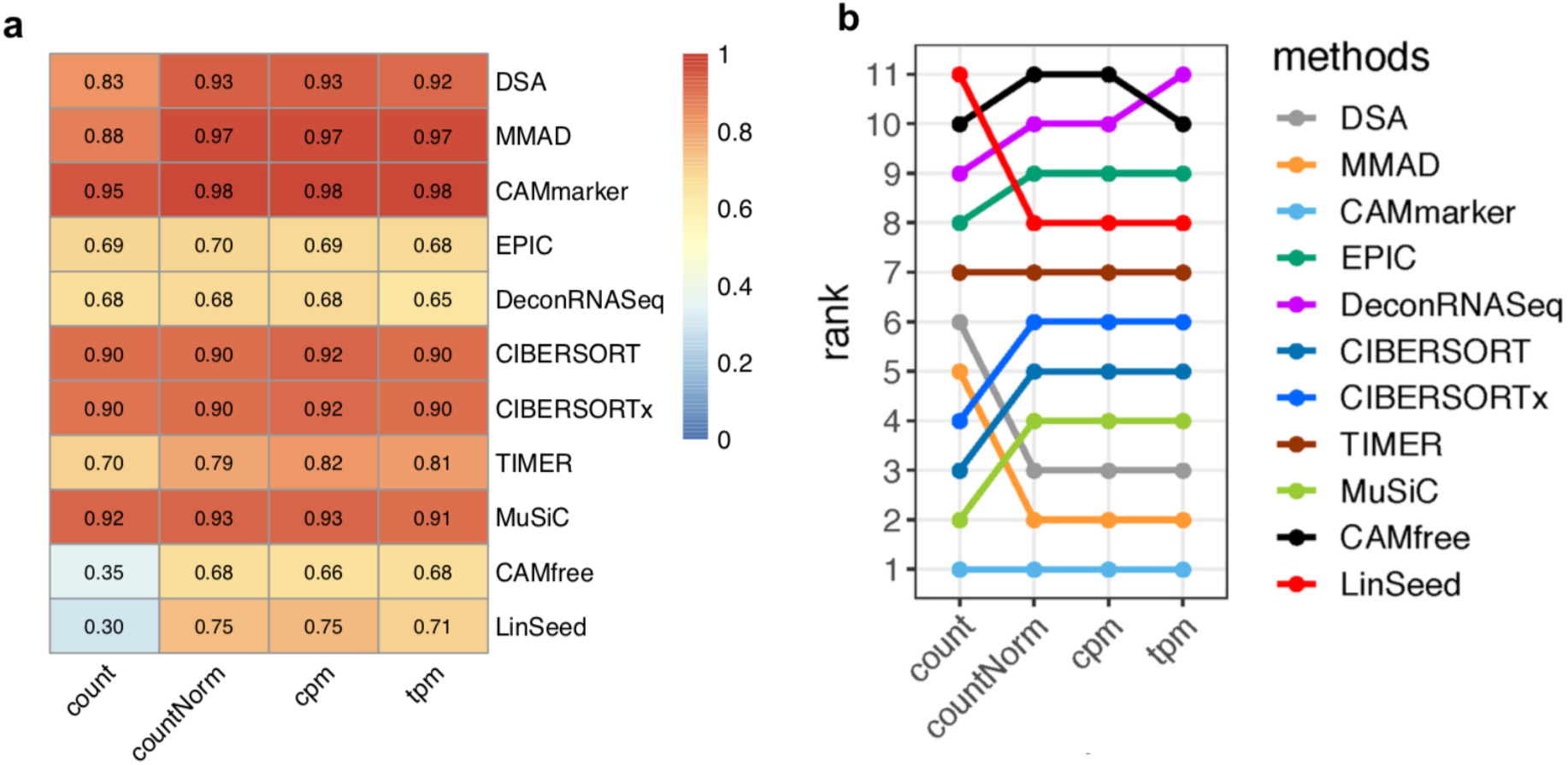
Evaluation results of Sim1_libSize. **a**, Heatmap of summarized evaluation results based on the Pearson’s correlation coefficients and **b**, rankings of tested deconvolution methods. In each heatmap, row indexes refer to the tested methods and column indexes refer to the quantification units (count, countNorm, cpm, and tpm).

We observed that three methods, CIBERSORT^7^, CIBERSORTx^8^, and MuSiC^16^, which implemented normalization procedures, showed decent performance (*r* ≥ 0.9, *mAD* ≤ 0.1) regardless of the selected quantification unit (Fig. 3a, Supplementary Fig. 11a). Six methods (DSA^11^, MMAD^12^, CAMmarker^13^, TIMER^10^, CAMfree^13^, and LinSeed^17^) showed improved accuracy after library size normalization (Fig. 3a, Supplementary Fig. 11a).

Contradicting to the Sim1_simModel (Supplementary Fig.3 and 4), we observed that the choice of quantification unit had a high impact on Sim1_libSize, which was reflected by discrepant rankings of tested methods (Supplementary Fig. 3b and 11b). As the only difference between the two benchmarking frameworks was the library size, we deduced that the inconsistent performance over different quantification units was due to the library size variation in the mixture dataset. We thus suggest researchers applying RNA-seq quantification units that are normalized by library sizes to mitigate the bias caused by varied library sizes unless indicated by the author of the method(MuSiC^16^) to use the count unit.

### Impact of cellular component number and weight matrix on deconvolution analysis

To investigate the joint impact of the cellular component number and weight matrix property, we designed the benchmarking framework Sim2 with six gradients of component number ranging from 5 to 10 and two types of weight matrices: ‘orthog’ and ‘real’ (Supplementary Fig. 2a and Supplementary Table 2 and 4). The ‘orthog’ weight matrix was generated by minimizing the condition number, and the ‘real’ weight matrix is constructed based on whole blood immune cell proportions in the real biological samples^21^(Methods). We discarded the CAMfree^13^ method in Sim2 due to the poor scalability of CAMfree^13^ on mixtures with large component numbers.

We found that nearly all deconvolution methods achieved higher accuracies with the ‘orthog’ weight matrices (Fig. 4a) than the ‘real’ weight matrices, indicating that the mathematical property of the weight matrix has a significant impact on deconvolution analysis. In the mixtures with five components (Comp 5), eight methods (DSA^11^, MMAD^12^, CAMmarker^13^, EPIC^14^, CIBERSORT^7^, CIBERSORTx^8^, MuSiC^16^, and LinSeed^17^) exhibited high accuracy levels(*r* ≥ 0.95, *mAD* ≤ 0.05) in the ‘orthog’ group (Fig. 4a and Supplementary Fig. 12a) while only three of those eight methods (CIBERSORT^7^, CIBERSORTx^8^, and MuSiC^16^) in the ‘real’ group achieved the same level of accuracy (Fig. 4b and Supplementary Fig. 12b).

**Fig.4.**
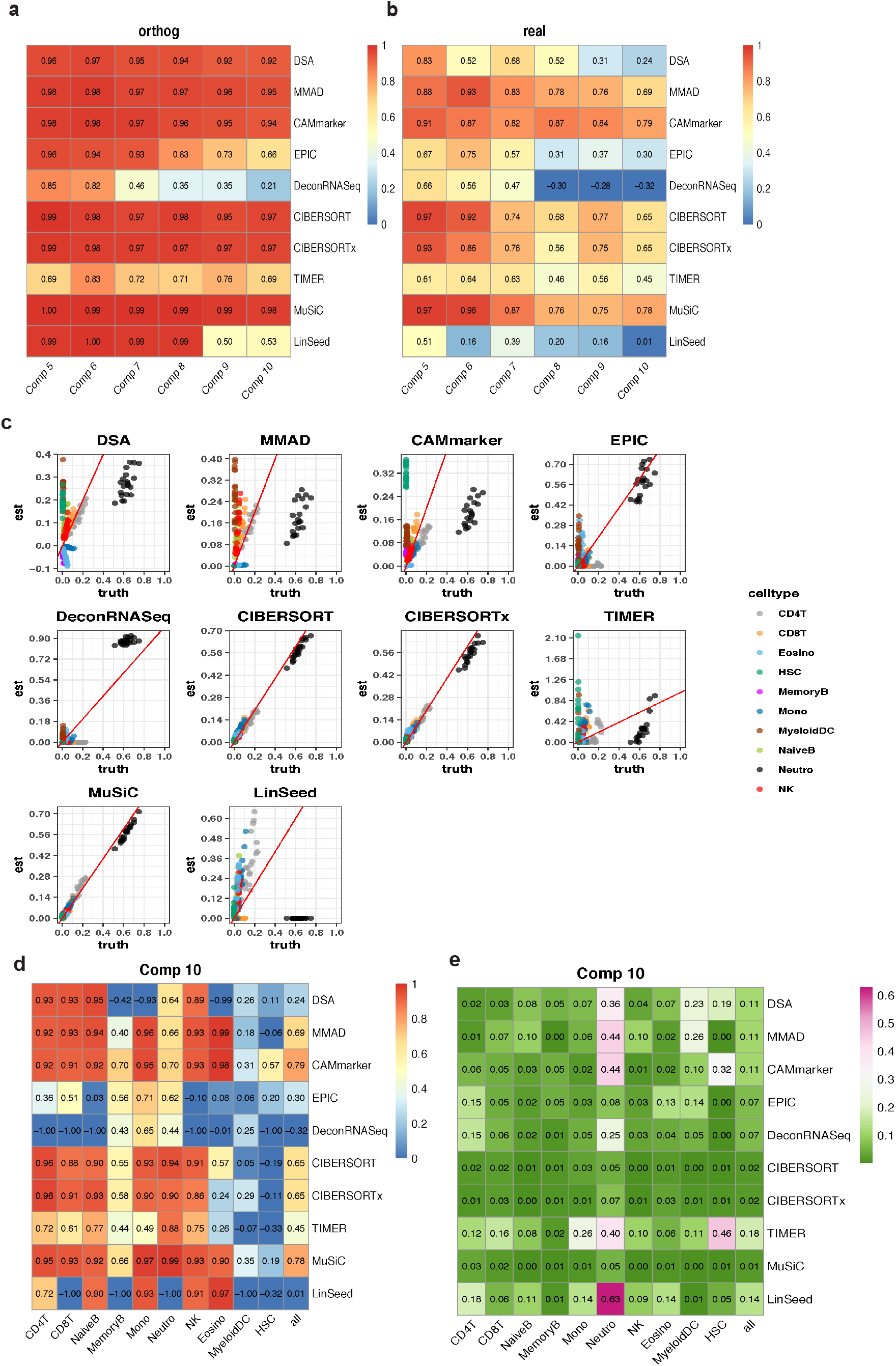
Evaluation results of Sim2. **a,b**, Heatmaps of summarized evaluation results based on the Pearson’s correlation coefficients with (**a**) ‘orthog’ weight matrix and (**b**) real weight matrix. In each heatmap, row indexes refer to the tested methods and column indexes refer to the cellular component numbers. **c**, Scatter plots of estimated weights vs. ground truths of mixtures with 10 cellular components. **d,e**, Cell-type specific evaluation metrics of mixtures consist of 10 cellular components based on (**d**) Pearson’s correlation coefficient and (**e**) Mean absolute deviance.

In addition to the impact of the weight matrix selection, cellular component numbers also affect deconvolution accuracy. In both ‘orthog’ and ‘real’ groups, the majority of methods exhibited poorer performance as cellular component number increasing (Fig. 4 a,b and Supplementary Fig. 12). It is also worth noting that none of the tested deconvolution methods showed a correlation larger than 0.9 with mixtures consist of large cellular component numbers (Comp 7 to Comp 10) in the ‘real’ group (Fig. 4b).

To further investigate the performance of deconvolution methods with large component numbers, we explored the accuracies of mixtures with 10 cellular components and the ‘real’ weight matrix by drawing scatters plots of estimations and ground truths (data corresponds to the last column of Fig. 4b and Supplementary Fig. 12b). Surprisingly, we found that the correlation evaluation metric, which was considered as the golden standard for the evaluation of deconvolution methods, cannot reflect the deviance of estimations from ground truths (Fig. 4c). However, the deviance of estimation can be reflected by another evaluation metric mAD (Supplementary Fig. 12). For instance, MMAD^12^ and CAMmarker^13^ performed relatively well on the correlation evaluation metric (*r* ≥ 0.65, Fig. 4b), but both methods had mAD values larger than 0.1, indicating large estimation deviance (Supplementary Fig. 12b). Consistent with the results from scatter plots (Fig. 4c), we found that the best performers were CIBERSORT^7^, CIBERSORTx^8^, and MuSiC^16^. All three methods achieved high accuracies on both correlation evaluation metric (*r* ≥ 0.65) (Supplementary Fig. 4b) and mAD evaluation metric (*mAD* ≤ 0.02) (Supplementary Fig. 12b) in the Comp 10 mixture with ‘real’ weight matrix.

To understand the impact of each cellular component on deconvolution analysis, we drew evaluation heatmaps with cell-type-specific correlation and mAD values (Supplementary Fig. 13, 14). Based on the evaluation heatmap of mixtures with ten cellular components and the ‘real’ weight matrix, which is the most complicated *in silico* mixture set in the Sim2 benchmark framework, we identified three best performers: CIBERSORT^7^, CIBEERSORTx^8^, and MuSiC^16^ (Fig. 4 d and e). First, we found that all three methods correctly estimated major cellular components (*r* ≥ 0.85, *mAD* ≤ 0.05), such as Neutrophils, CD4T, and CD8T in the respective mixtures. Second, while all three methods failed to estimate the linear trend of proportions of rare cell subpopulations that occupies less than 1% in the mixture, such as Myeloid DC and HSC (Hematopoietic Stem Cells) (*r*: −0.19 ~ 0.35), they correctly identified them as minor components and did not attribute the percentages of other cell types to these rare cell populations (*mAD*: 0 ~ 0.01). Moreover, because none of the tested deconvolution methods showed good accuracies in both correlation and mAD metrics with Myeloid DC and HSC (Figure 4 d and e), we concluded that none of the currently developed deconvolution methods could not reliably estimate some rare cellular populations that have proportions less than 1%. Finally, we also discovered that marker-gene based methods like DSA^11^, MMAD^12^, and CAMmarker^13^ showed high mAD values (Figure 4d and e), indicating larger deviances in their estimations in the major components(*mAD*: 0.36 ~ 0.44)(Fig. 4e).

By inspecting cell-type-specific evaluation results of ‘real’ weight matrices across 6 component gradients, we found that introducing rare cellular components MyeloidDC in the Comp 7 mixture caused the deterioration of deconvolution performance, which might be due to the close relationship between MyeloidDC to the monocytes^22^. However, introducing relatively distinct HSC in the Comp 8 mixture further exacerbated the performance deterioration (Supplementary Figures 13 and 14, ‘real’ group). Therefore, we concluded that the deterioration of deconvolution performance on mixtures with large component number is due to the confounding effect of the highly correlated cellular component and the rare cellular component in the mixture dataset.

### Impact of tumor content on deconvolution analysis

Unknown biological content, such as tumor content, is another major factor that influences deconvolution analysis for several reasons. First, unknown content could be treated as a source of noise unless explicitly modeled by deconvolution methods^7,14^. Second, unknown content is not counted in the estimated cell-type proportions and violates the sum-to-one assumption applied by the majority of deconvolution methods^2,9^.

To study the impact of unknown biological content on deconvolution analysis, we designed a benchmarking framework that contains mixtures with three sets of tumor spike-ins: the ‘small’ group refers to mixtures with low levels of tumor spike-ins (0 - 20%), the ‘large’ group refers to mixtures with high levels of tumor spike-ins (70 - 90%), and the ‘mosaic’ group refers to mixtures with more dynamic levels of tumor spike-ins (5% - 95%). Tumor spike-ins were introduced to the 12 mixture sets generated in the Sim2 framework to analyze the joint impact of the component numbers, weight matrix properties, and unknown biological contents (Supplementary Fig. 2b, Methods). In the performance assessment step, we used two sets of ground truths to derive evaluation results that represent different measurement scales (Supplementary Table 5, Methods). The first set of ground truths used the absolute proportions of immune cell types and led to ‘absolute’ deconvolution accuracy. The second set of ground truths used the relative proportions of immune cells and led to ‘relative’ deconvolution accuracy. In this set of analyses, we considered additional settings of deconvolution methods that were relevant to the tumor content. Thus, we evaluated eleven methods and two specific method settings TIMERtumor^10^ and EPICabsolute^14^, which are tailored for deconvolution analysis with unknown tumor contents (Methods, Supplementary Table 3).

Our results indicated the weight matrix property as the leading factor that affected deconvolution accuracy because the ‘orthog’ group presented higher accuracies throughout all deconvolution methods and tumor content conditions (Fig. 5a, b and Supplementary Fig. 15). In addition to the weight matrix property, we found that the size of tumor content also affected deconvolution accuracy as we observed deconvolution methods performed better on mixtures with smaller tumor content (Fig. 5a, b and Supplementary Fig. 15). Moreover, we found that all methods showed inconsistent performance with the ‘mosaic’ mixture group when evaluated on different measurement scales (Fig. 5a, b and Supplementary Fig. 15). For instance, in the ‘mosaic’ column, CIBERSORT^7^ and CIBERSORTx^8^ showed higher accuracies (*r*: 0.69~0.95, *mAD*: 0.03) in the relative measurement scale (Fig. 5a and Supplementary Fig. 15a) than in the absolute measurement scale (*r*:0.4~0.97, *mAD*: 0.06~0.07) (Fig. 5b and Supplementary Fig. 15b). Methods like DSA^11^, MMAD^12^, CAMmarker^13^, EPIC^14^, EPICabsolute^14^, TIMER^10^, TIMERtumor^10^, and MuSiC^16^ showed higher accuracies in the absolute measurement scale (*r*: 0.33 ~ 0.9, *mAD*: 0.21) (Fig. 5b and Supplementary Fig. 15b) than in the relative measurement scale (*r*: 0.22 ~ 0.68, *mAD*: 0.17) in the ‘mosaic’ column(Fig. 5a and Supplementary Fig. 15a).

**Fig.5.**
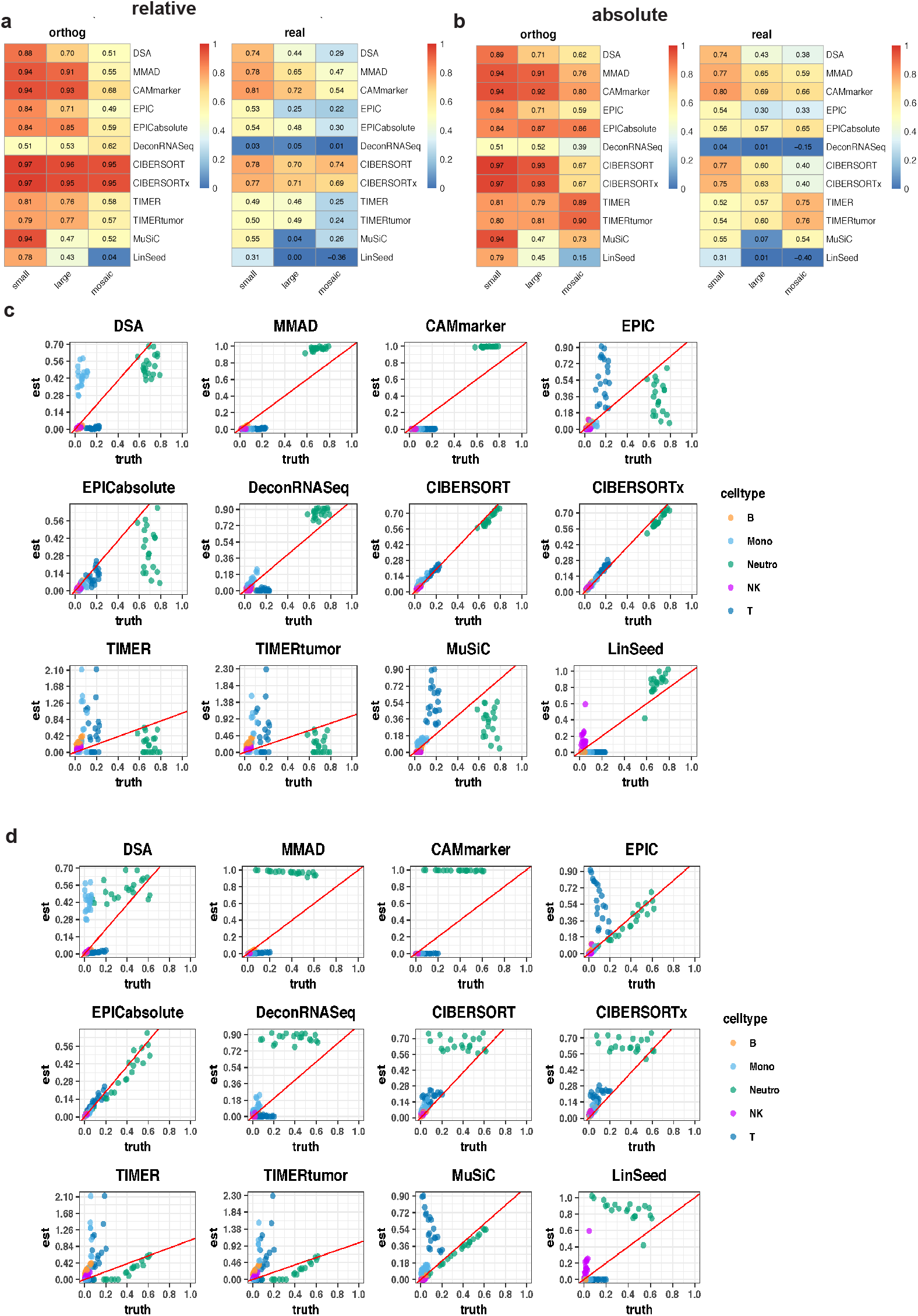
Evaluation results of Sim3. **a,b**, Heatmaps of summarized evaluation metric based on Pearson’s correlation coefficients on the (**a**) relative measurement scale and (**b**) absolute measurement scale. In each heatmap, row indexes refer to the tested methods and column indexes refer to the types of tumor spike-ins (small, large, and mosaic). **c,d**, Scatter plots of estimated weights vs. ground truths of mixtures consist of 5 cellular components and mosaic tumor spike-ins. (**c**) estimated weights vs. relative ground truth (**d**) estimated weights vs. absolute ground truth.

To further investigate the performance of deconvolution methods under the cell-type resolution, we drew scatter plots of estimations from 5 Comp mixtures with ‘mosaic’ tumor spike-ins and ‘real’ weight matrix (Fig. 5 c,d). In the relative measurement scale, CIBERSORT^7^ and CIBERSORTx^8^ were the top performers and achieved high accuracy (*r* ≥ 0.95, *mAD* ≤ 0.05) (Fig. 5c and Supplementary Fig. 16). However, in the absolute measurement scale, EPICabsolute^14^ was the top performer and correctly estimated the absolute immune cell proportions (*r* ≥ 0.95, *mAD* ≤ 0.05) (Fig. 5d and Supplementary Fig. 17). Based on inconsistent evaluation results from two measurement scales, we suggest researchers pay attention to the impact of measurement scales when performing deconvolution analysis on mixtures with unknown contents.

Next, we checked the robustness of the three best performers in terms of component number and tumor content in the ‘real’ weight matrix group. The robustness of CIBERSORT^7^ and CIBERSORTx^8^’s performance to the component number is high on the mAD evaluation metric (*mAD*: 0.02 ~ 0.05) in the relative measurement scale (Supplementary Fig. 16b). EPICabsolute^14^ also showed high robustness to the component number on the mAD evaluation metric (*mAD*: 0.02 ~ 0.07) in the absolute measurement scale(Supplementary Fig. 17b). We found that having a larger variance in tumor content will increase the accuracy of EPICabsolute^14^, as we observed that with mosaic tumor spike-ins, EPICabsolute achieved higher accuracies (*r*: 0.31~0.95, *mAD*: 0.02~0.05) than other tumor spike-in groups(*r*: 0.17~0.84, *mAD*: 0.02~0.07) (Supplementary Fig. 17) in the absolute scale. Consistent with the observation in Sim2, we observed decreasing accuracies of CIBERSORT^7^, CIBERSORTx^8^, and EPICabsolute^14^ with the increasing component number (Supplementary Fig. 16a and Supplementary Fig. 17a), and we deduced this phenomenon is due to the difficulty of current deconvolution methods estimating rare subpopulations and closely related cell-types.

Our results revealed the impact of unknown biological content on deconvolution analysis. We found both size (large vs. small spike-ins) and variance (large vs. mosaic spike-ins) of unknown content affected deconvolution analysis. We also observed a discrepancy in performance evaluation when used different measurement scales. In the relative scale, we concluded CIBERSORT^7^ and CIBERSORTx^8^ were the top performers, while in the absolute scale, EPICabsolute^14^ was the top performer.

## Discussion

In this study, we designed three *in silico* benchmarking frameworks to systematically explore the impact of several biological and technical factors. We identified top-performing deconvolution methods for each framework and clearly illustrated the strengths and limits of these tested methods under different application scenarios. Moreover, we offered several strategies to mitigate systematic biases caused by different technical and biological factors such as varied library sizes, simulation models, and cellular compositions.

In the first framework (Sim1), we explored the impact of noise structure under different noise levels. We identified CAMmarker, MMAD, DSA, and CIBERSORT as the best performers since these methods showed high accuracy and high robustness to diverse noise levels. For the noise structure, we identified the negative binomial as the best simulation model that captures the essential characteristics of real data. In the second framework (Sim2), we explored the impact of the cellular component number and the weight matrix property. We identified CIBERSORT, CIBERSORTx, and MuSiC as top-performers since these two methods achieved high accuracies across a gradient of cellular component numbers with both ‘orthog’ and ‘real’ weight matrices. We also found all marker-gene based methods exhibited larger estimation deviances from ground truths, this type of estimation biases is reflected in the scatter plots and can be quantitatively captured by the mAD evaluation metric, indicating the necessity of using mAD as an auxiliary evaluation metric for deconvolution performance assessment. In the third framework (Sim3), we explored the impact of unknown biological content and measurement scales. In the relative measurement scale, CIBERSORT and CIBERSORTx were the best performers. In the absolute measurement scale, EPICabsolute was the best performer. Our analysis also illustrated different evaluation results under the absolute and relative measurement scale, which have been overlooked in the previous deconvolution benchmarks.

Based on the observations in this benchmark, we give the following suggestions for best practices of deconvolution analysis and evaluations. For the *in silico* benchmarking data generation, we suggest researchers 1) Use the negative binomial model as the primary simulation model for *in silico* mixture data generation. 2) Referencing real biological composition data when building weight matrices. 3) Consider at least two evaluation metrics. One is used for checking linear concordance between estimation and ground truth, and the other one is used for checking estimation deviances. 4) In the context of unknown biological content, beware of the influence caused by different measurement scales(absolute vs. relative). 5) Constructing multi-factor conditions on a large scale to ensure the robustness and comprehensiveness of the benchmark.

For deconvolution analysis, we suggest researchers 1) Use the quantification unit (countNorm, cpm, or tpm) that is normalized by library sizes. 2) Check for the compositional information from previous publications. When the targeted tissue type has a relatively stable composition over several samples, consider using deconvolution methods that are robust to non-orthog weight matrices such as CIBERSORT, CIBERSORTx, and MuSiC. When an unknown cellular component is expected (i.e., tumor sample) and the researcher needs to derive absolute proportion, consider methods like EPIC, which is specifically tailored for deconvolution with unknown content. 3) When referencing benchmark paper to select the optimal method, beware of different technical factors that might derive different estimation accuracies such as the resolution of analysis(number of cellular components), the variance of proportions across samples(weight matrix property), reference selection, evaluation metric selection, and measurement scale selection.

In addition to the suggestions mentioned above, previous benchmark publications also clarified the impact of signature matrices^1^, multicollinearity issue^7^, spill-over effects^3,23^ caused by missing cellular components in the reference, minimal detection fraction^3^, background predictions^3^, marker/signature gene selection^4,6^, the variance between reference and mixture sources^4^. Some deconvolution methods like CIBERSORT, CIBERSORTx, and MuSiC can derive both cell-type-specific expression and composition signals. However, by far, all independent deconvolution benchmark studies have been focused on the accuracy of compositional information^3,6^. More benchmarks that derive accuracies of cell-type-specific expression estimation are still in need.

For the future advancement of deconvolution analysis on RNA-seq data, we suggest more efforts be put into the refinement of simulation models to generate more authentic *in silico* testing environments that mimic diverse application scenarios. The weight matrix property was revealed as the most important factor affecting deconvolution analysis in this study and have been overlooked by the community. Therefore, more studies on the cellular compositional information and its corresponding effects on deconvolution analysis are still in need. Devotions on improving in silico benchmark generation strategy could further enhance the efficiency of deconvolution method development and enable a wide range of clinical applications.

## Methods

### Data processing

Raw SRA files were downloaded from the GEO repository, processed by SRA Toolkit (2.10.0)^24^, and reads were aligned to the hard masked human reference GRCh38 (v95) using alignment tool STAR (2.6.1)^25^, and quantification was performed with RSEM (1.3.1)^26^ with default parameter settings. Quantification matrices with the count, tpm, and fpkm units were loaded into R (3.6.1)^27^ for feature ID transformation, duplication removal, and low-abundant gene removal. For low-abundant gene removal, we relied on two parameters: minimum sample threshold (GSE113590^28^ - 4, other datasets - 5) and minimum expression threshold (10 counts, 1 tpm, and 1fpkm). For instance, the filtering parameter (5, 10) is used to retain genes with more than 10 counts in at least 5 samples. GSE113590 only has 4 samples per cellular category, and we set the minimum sample thresholds as 4. In the Sim1, we performed filtering independently on each dataset with a minimum sample threshold set at 5. For Sim2 and Sim3, we first concatenated samples into one matrix and then performed filtering with a minimum sample threshold set at 10. For the information of datasets involved in Sim1, Sim2, and Sim3, please refer to Supplementary Table 4.

### Marker gene selection

For the marker gene selection, we selected genes that are highly expressed in the targeted cell-type and lowly expressed in other cell-types. The expression threshold is set at the 80th percentile for high expression (the targeted group) and 50th percentile for low expression (other groups). Ideally, it would be nice if all samples pass the criteria; however, to successfully derive marker genes with a larger number of cellular components, we gradually relaxed the threshold (the percentage of samples pass the criteria, initial value p = 0.95) by a step parameter (default value s = 0.03) until there are at least two marker genes determined.

### Signature gene selection

We performed differential expression testing on all cell-type pairs (all combinations of 2 elements) using DESeq2^29^. Then we selected genes with *p_adj_* ≤ 0.01 and *log2FoldChange* ≥ 10.

### Benchmarking framework construction

Three benchmarking frameworks are constructed to study the impact of different technical and biological factors on deconvolution analysis (Figure 1). We created simulated mixture data M (N by J) by multiplying signature gene profiles S (N by K) to the predefined weight matrix W (K by J). Here, N is the number of genes, J is the number of samples, and K is the number of cellular components. The noise term *ε* is used to model sample to sample variability where the value of *ε* determines the noise level.

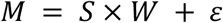

#### Sim1

In the Sim1, we aimed at understanding the impact of noise from different aspects such as noise structure and noise level. Sim1 consists of two sub frameworks: Sim1_simModel and Sim1_libSize, where Sim1_simModel focuses on the noise structure, and Sim1_libSize focuses on noise caused by varied library sizes.

#### Sim1_simModel

In this benchmarking framework, we mainly focused on the impact of the simulation model that was used to generate noise. We selected three models for this study, which are the normal, log-normal, and negative binomial models. For each simulation model, we generated ten levels of noise where the magnitude of the noise is controlled by a corresponding variance term in each model.

#### Normal model

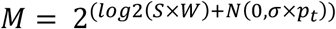

#### Log-normal model

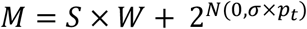

In both Log-normal and Normal simulation models, the level of noise is controlled by the product of a constant variance parameter *σ* and a perturbation level parameter *p_t_*. In this study, we set *σ* to 10 based on previous publications^7^ and set *p_t_* as a length-10-vector (0, 0.1, 0.2, …, 0.9).

#### Negative binomial model

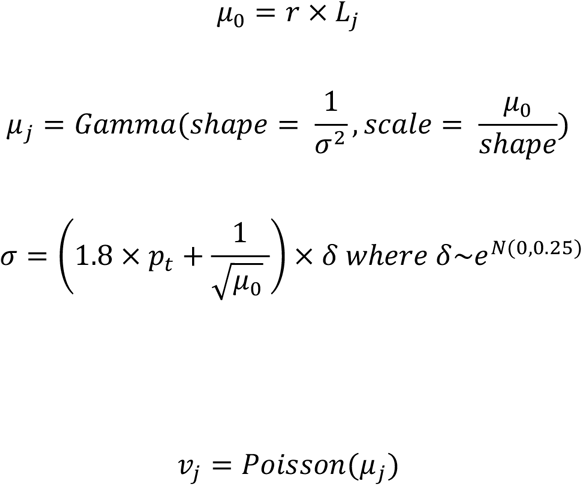

We followed the simulation process suggested by Law *et al*.^19^ and used *p_t_* to control the noise level for simulation. r is a vector of genomic feature proportions, *L_j_* is the library size and, *μ*_0_ is the expected gene expression in the simulation. In the negative binomial model, two layers of variance are added from the Gamma distribution and Poisson distribution. We derived sample gene expression vector *μ_j_* from Gamma sampling to model biological variance. In the Gamma distribution, the variance is determined by shape parameter *σ*. We used *p_t_*, a length-10 vector (0.1, 0.2, …, 0.9, 1), to regulate the value of *σ* to control the noise level in the negative binomial simulation. Then we performed Poisson sampling to model technical variance and get the final simulated expression vector.

To ensure the universality of our conclusion on different datasets, we applied the Sim1 framework on 3 blood datasets to generate reference and *in silico* mixtures (Supplementary Fig.1). Different from previous studies that concatenate samples derived from different datasets, we generated 3 sets of simulated mixtures and 3 sets of references independently. And then used combinations of mixtures and references to generate 9 replicated testing environments for each noise level. For one testing environment, there are 9 (3 times 3) deconvolution results from which 6 of them have mixture-reference pairs derived from different sources. For simplicity, we only presented the averaged performance across 9 mixture-reference pairs, but the impact of mixture-reference variance is considered in this analysis. Above mentioned mixture-reference variance modeled in Sim1 is named as other noise sources in Supplementary Table 2.

To understand the impact of quantification units over different application scenarios, we generated simulations of the most commonly used RNA-seq quantification units: count, countNorm, cpm, and tpm.

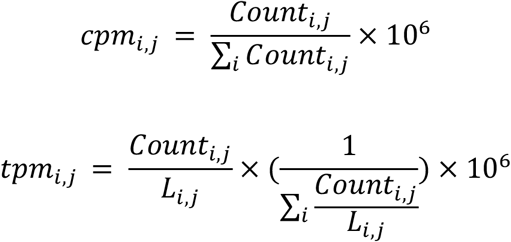

Here j is the index of the sample and i is the index of the gene. cpm is normalized by library size. countNorm is acquired from cpm units with every value rounded to the integer. tpm is normalized by both library size and feature-length.

#### Sim1_libSize

In this testing framework, we mainly focused on bias derived from varied library sizes. We first simulated mixtures based on the negative binomial model with the lowest level of noise in Sim1_simModel (*p*_1_ perturbation level). The library size variation is controlled by the library size parameter *L_j_* in the negative binomial model. For every simulation dataset that consists of 20 simulated profiles, we set the library size of the first ten samples as 12 million reads and the remaining ten samples as 24 million reads (Supplementary Fig. 1b).

#### Sim2

In this benchmarking framework, we studied the impact of cellular component numbers and the mathematical property of the weight matrix (Supplementary Fig.2a). Mixtures are generated based on the negative binomial model with the *p_1_* level noise. For component number, we generated six sets of mixtures from 5 components up to 10 components. For the weight matrix, we generated two sets of weight matrix: orthog and real.

#### Weight simulations

**‘Orthog’** refers to the idealized weight matrix with a small condition number, which provides a relatively optimal mathematical condition for deconvolution analysis. We first simulated 1000 matrices (K by J) by randomly sampling weights from a uniform distribution and then rescaled sampled weights so that for each mixture sample, all components sum to 1. Among 1000 proportion matrices, we picked the one weight matrix that has the smallest condition number. **‘Real’** refers to the weight matrix that mimics immune cell compositions in the real whole blood sample. We generated weights based on uniform distribution with min and max value defined based on previous observations of whole blood samples^21^ and then rescaled weights so that all components sum to 1.

#### Sim3

In this benchmarking framework, we studied the impact of unknown biological content and measurement scales (Supplementary Fig.2b). To study unknown biological content, we generated mixtures with tumor content spike-ins. In total, we created three sets of tumor spike-ins: small, large, and mosaic. Tumor proportions are sampled from uniform distributions and only differ in parameters used to set minimum and maximum values in the sampling. ‘Small’ tumor spike-ins are sampled within the range 0-0.2, ‘large’ tumor spike-ins are sampled within the range 0.7-0.9, and ‘mosaic’ tumor spike-ins are sampled within the range 0.05-0.95. We then added three sets of tumor spike-in proportions to the weight matrices generated in the Sim2 and rescaled them to have proportions of all components sum to 1. After defining weights, we performed *in silico* mixing in the count unit and then normalized it to other quantification units. To study the impact of the measurement scale, we generated two sets of evaluations where one used absolute proportions of immune components as the ground truth and the other used relative proportions of immune components as the ground truth. The toy example of the absolute measurement scale and the relative measurement scale is in Supplementary Table 5.

#### Assessment of deconvolution performance

J is the total number of mixture samples in a dataset and j is the sample index. *x_j_* is the estimated proportion of sample j and *y_j_* is the ground truth of sample j. When a deconvolution returns NA values, we directly assign highest penalty for the evaluation metrics: *r* = −1, and *mAD* = 1.

### Pearson Correlation Coefficient (r)

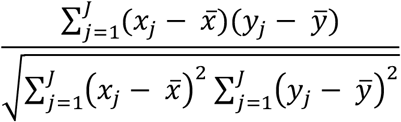

### Mean Absolute Deviance (mAD)

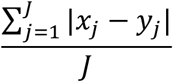

### Datasets description

1. **GSE60424**^30^ **-** Consists of 134 RNA-seq profiles of 6 immune cell types and whole blood from both healthy donors and donors with five immune-associated diseases.
2. **GSE113590**^28^ **-** Consists of 32 CD8 T cell RNA-seq profiles from peripheral blood, colorectal tumor samples, and lung tumor samples.
3. **GSE64655**^31^ **-** Consists of 56 RNA-seq profiles of 6 immune cell types and peripheral blood from two vaccinated donors.
4. **GSE51984**^32^ - Consists of 24 RNA-seq profiles of 5 immune cell types and total white blood cells from healthy donors
5. **GSE115736**^33^ **-** Consists of 42 RNA-seq profiles of 12 immune cell types from healthy donors.
6. **GSE118490**^34^ **-** HCT116 profiles (unknown tumor content in Sim3)

## Data and code availability

All data and codes are available in the https://github.com/LiuzLab/paper_deconvBenchmark under MIT license.

## Author contributions

H. J. designed, planned, and conducted data analysis and wrote the manuscript. Z.L. supervised the analysis and wrote the manuscript. All authors read and approved the final manuscript.

