## Supplementary Files for "A comparative study of deconvolution methods for RNA-seq data under a dynamic testing landscape"

**Supplementary Table 1: Three benchmarking frameworks commonly applied**

| Framework type | Sample source | Ground truth | Flexibility | Cost-effectiveness | Similarity to the biological condition |
|---|---|---|---|---|---|
| **in silico** | simulation | yes | High | High | Low |
| **in vitro** | experimentally generated mixture samples | yes | Medium | Medium | Medium |
| **in vivo** | Samples obtained directly from a biological organism | Usually no | Low | Low | High |

### Supplementary Table 2: Summary of *in silico* benchmarking frameworks in this study

| **Benchmarking frameworks** | **Factor 1** | **Factor 2** | **Factor 3** | **Factor 4** | **Factor 5** | **Number of conditions** | **Dataset index** |
|---|---|---|---|---|---|---|---|
| **Sim1_simModel** | Simulation model | Noise level | Other noise sources | Units | | 1080 | 1,3 and 4 |
| **Sim1_libSize** | Units | Noise level | | | | 360 | 1,3 and 4 |
| **Sim2** | Weight matrix | Component number | Units | | | 48 | 1,3,5 and 5 |
| **Sim3** | Tumor content | Measurement scale | Weight matrix | Component number | Units | 288 | 1,3,4,5 and 6 |
| Total number of conditions: 1, 776<br><br>**Summary of factors:**<br><br>Simulation model: 3 (normal, log-normal, and nb)<br><br>Noise level: 10 (P1-P10)<br><br>Units: 4 (count, countNorm, cpm, and tpm)<br><br>Weight matrix: 2 (orthog and real)<br><br>Component number: 6 (5 - 10)<br><br>Tumor content: 3 (small, large, and mosaic)<br><br>Measurement scale: 2 (relative and absolute)<br><br>Other noise sources: 9 (variance between reference and mixture) | | | | | | | Data index corresponds to the index in the dataset description |

**Supplementary Table 3: Summary of tested methods in this study**

| Deconvolution methods | Description | Type |
|---|---|---|
| DSA[11] | Least squares or quadratic programming | Marker-based |
| MMAD[12] | Maximum likelihood | Marker-based |
| CAMmarker[13] | Simplex approach | Marker-based |
| CIBERSORT[7] | Support vector regression | Reference-based |
| EPIC[14] | Weighted least squares | Reference-based |
| TIMER[10] | Least-squares | Reference-based |
| DeconRNASeq[15] | Least-squares | Reference-based |
| MuSiC[16] | Iterative weighted least squares | Reference-based |
| LinSeed[17] | Simplex approach | Reference-free |
| CAMfree[13] | Simplex approach | Reference-free |
| Additional information:<br><br>We had to make some modifications to the name of some methods since we used different parameter settings for the same method.<br><br>1. CAMmarker and CAMfree are all derived from R package CAMTHC[13]. We named the marker-based approach CAMmarker and the reference-free approach CAMfree.<br>2. We referred EPIC[14] with the unknown content estimation as EPICabsolute.<br>3. We referred TIMER[10] with the additional filtering process specific for tumor genes as TIMERtumor. | | |

**Supplementary Table 4: Cell types and datasets involved in each analysis**

| Analysis | Relevant Figures | Cell-types | Datasets |
|---|---|---|---|
| Variance Analysis | Fig 2 c-h, SuppFig 6 - 10 | CD8 T cells<br>Whole Blood,<br>Simulated Mixtures (T, B and Mono) | GSE113590<br>GSE60424<br>GSE51984 |
| Sim1_simModel | Fig 2 a,b, SuppFig 3 and 4 | Simulated Mixtures (T, B and Mono) | GSE60424<br>GSE51984<br>GSE64655 |
| Sim1_libSize | Fig 3, SuppFig 5 | Simulated Mixtures (T, B and Mono) | |
| Sim2 | Fig 4, SuppFig 12 and 13 | 6 gradients of cell-types | GSE60424<br>GSE51984<br>GSE64655<br>GSE115736 |
| Sim3 | Fig 5, SuppFig 15, 16 and 17 | 6 gradients of cell-types, HCT116 | GSE60424<br>GSE51984<br>GSE64655<br>GSE115736<br>GSE118490 |
| Additional information of 6 gradients of cell types:<br>Comp6 - T, B, Monocytes, Neutrophils, and NK cells<br>Comp7 - T, B, Monocytes, Neutrophils, NK cells, and Eosinophils<br>Comp8 - T, B, Monocytes, Neutrophils, NK cells, Eosinophils, and Myeloid DC<br>Comp9 - CD4 T, CD8 T, B, Monocytes, Neutrophils, NK cells, Eosinophils, Myeloid DC and CD34+ HSC<br>Comp10 - CD4 T, CD8 T, Naïve B, Memory B, Monocytes, Neutrophils, NK cells, Eosinophils, Myeloid DC and CD34+ HSC | | | |

**Supplementary Table 5: Toy example of different measurement scales**

| | T cell | B cell | Unknown contents | Sum |
|---|---|---|---|---|
| Absolute scale weights | 0.3 | 0.2 | 0.5 | 1 |
| Relative scale weights | 0.3/(0.5) = 0.6 | 0.2/(0.5) = 0.4 | / | 1 |

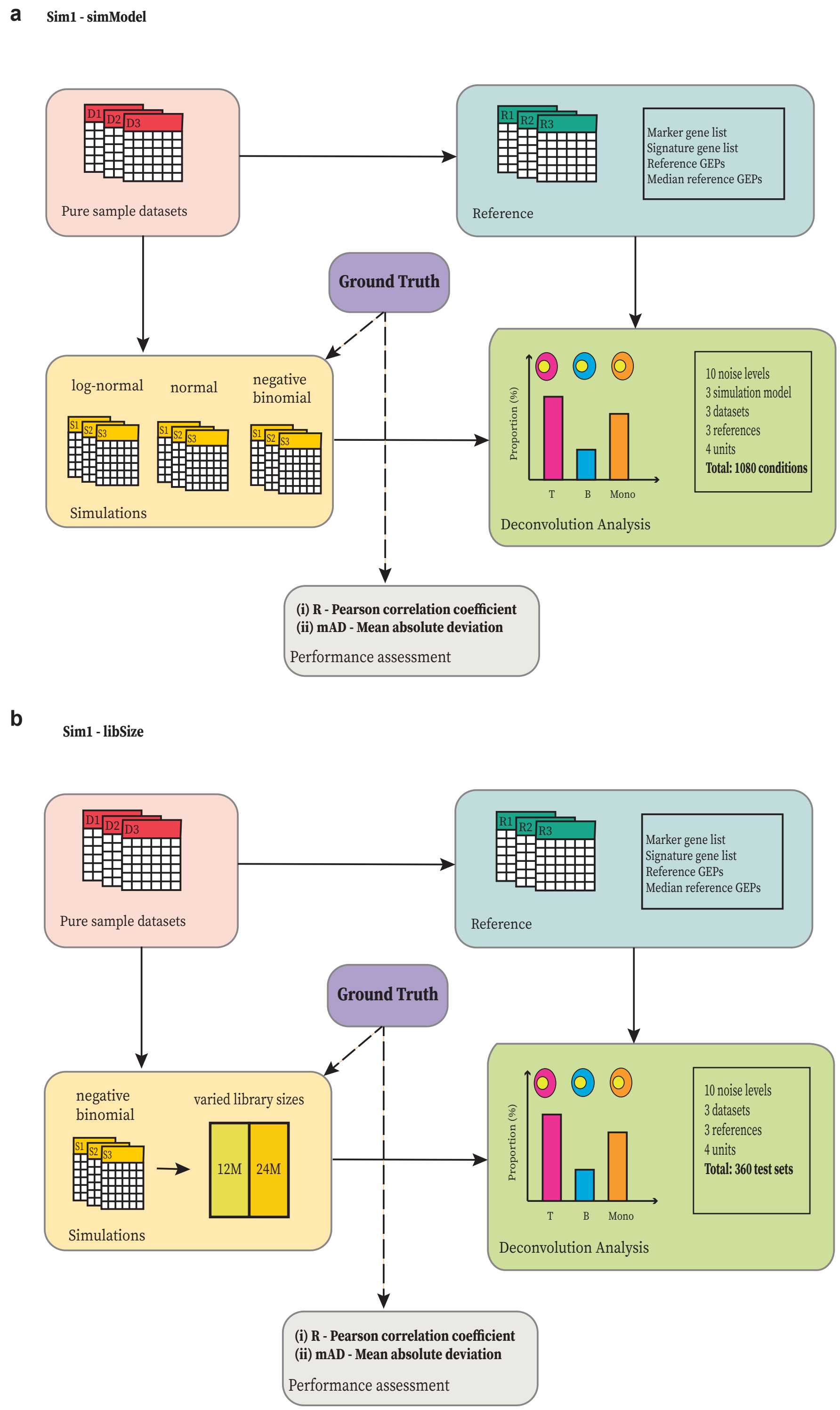


**Supplementary Fig. 1| Outline of benchmarking framework Sim1**

**a,** Outline of benchmarking framework Sim1_simModel. **b,** Outline of benchmarking framework Sim1_libSize

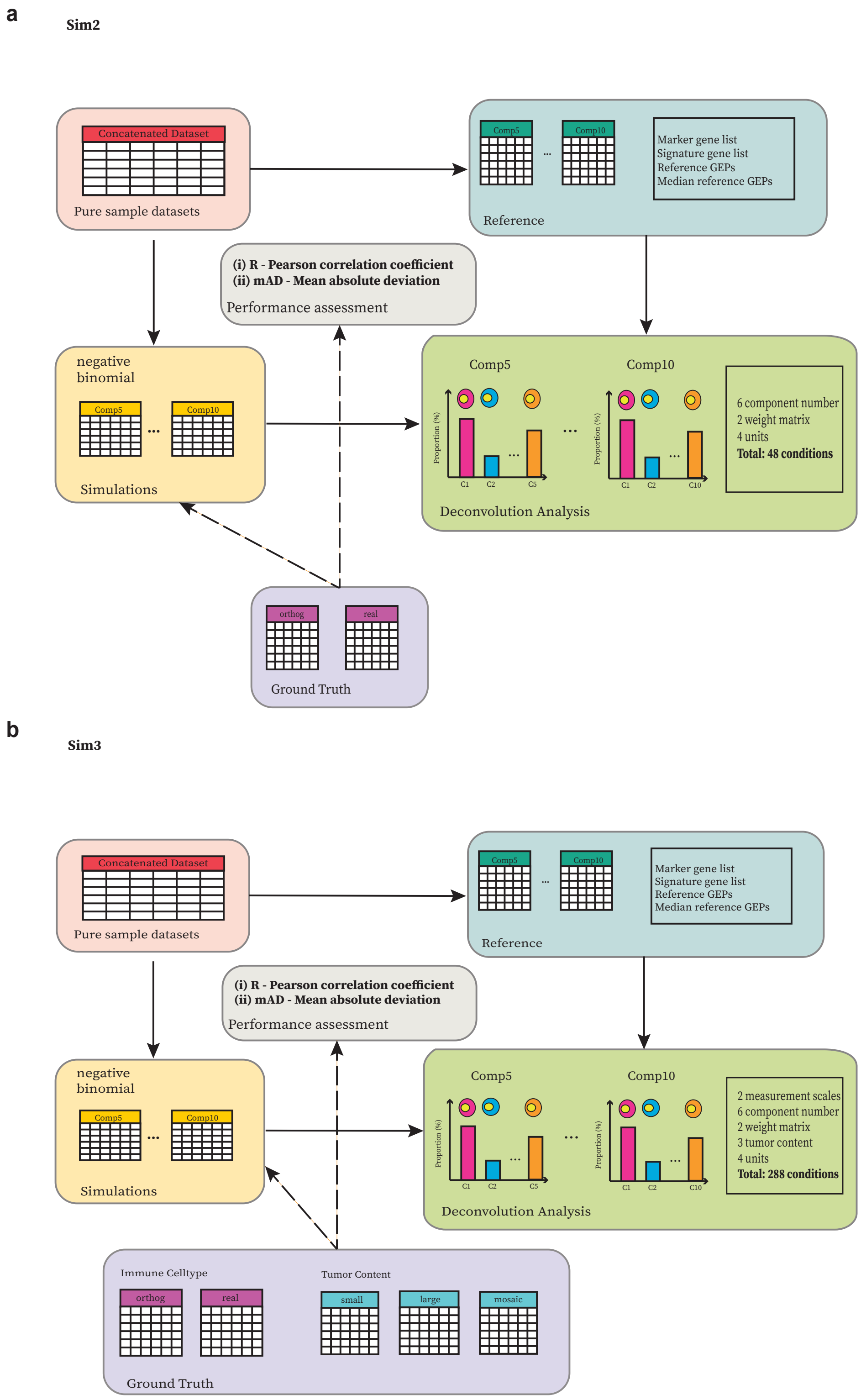


**Supplementary Fig. 2| Outline of benchmark frameworks Sim2 and Sim3**

**a,** Outline of benchmarking framework Sim2. **b,** Outline of benchmarking framework Sim3.

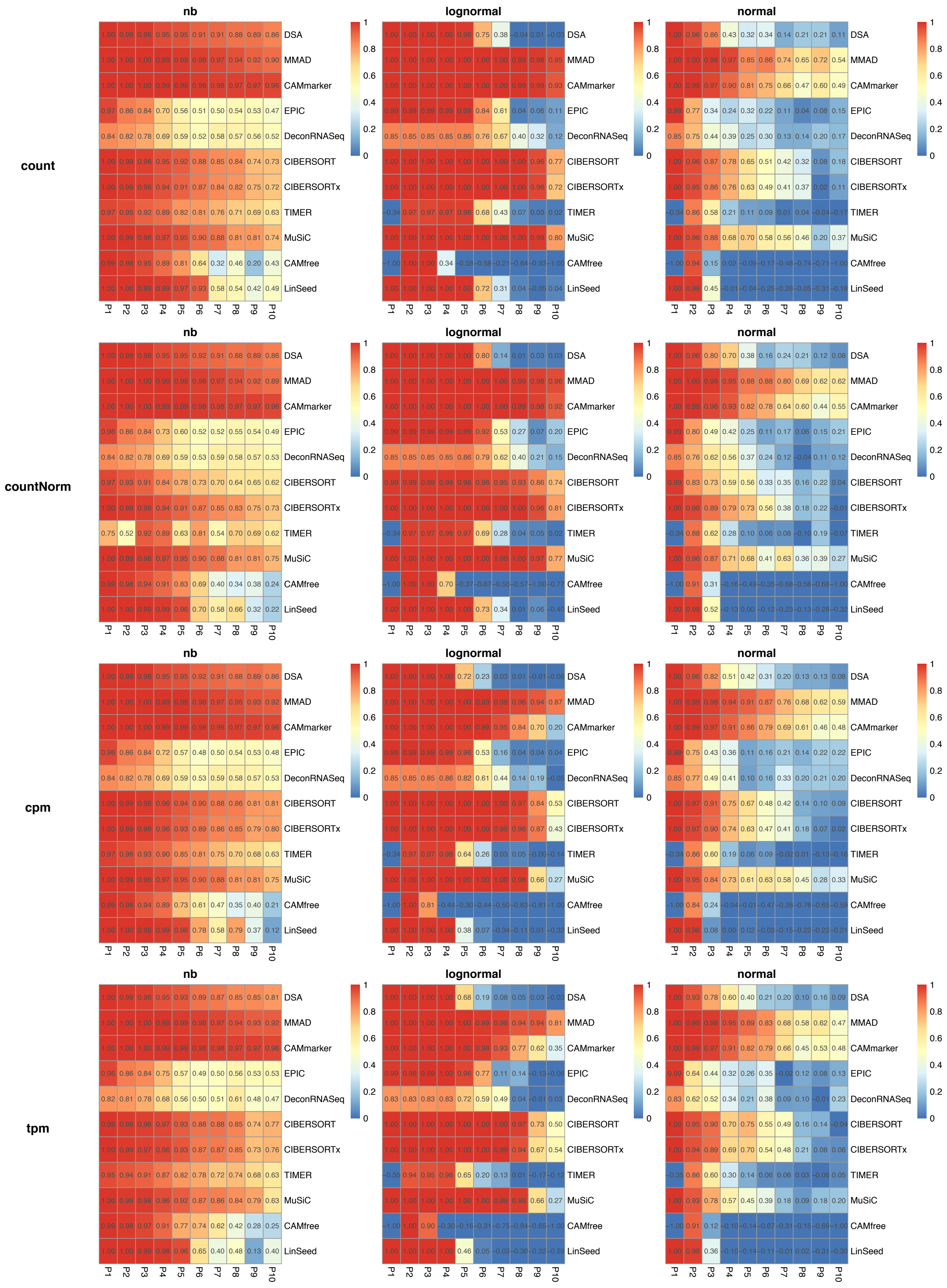
count
countNorm
cpm
tpm
nb
lognormal
normal
DSA
MMAD
CAMmarker
EPIC
DeconRNASeq
CIBERSORT
CIBERSORTx
TIMER
MuSiC
CAMfree
LinSeed
P1
P2
P3
P4
P5
P6
P7
P8
P9
P10
1
0.8
0.6
0.4
0.2
0

### Supplementary Fig. 3| Correlation heatmaps of sim1_simModel

The deconvolution results of Sim1_simModel are organized by 12 heatmaps where each row panel refers to the results derived from the same quantification units and each column panel refers to the results derived from the same simulation model. In each heatmap, row indexes refer to the tested deconvolution methods and column indexes refer to the noise level. Each cell in the heatmap indicates averaged correlations of 9 testing sets (3 mixture sets × 3 reference sets)(Methods).

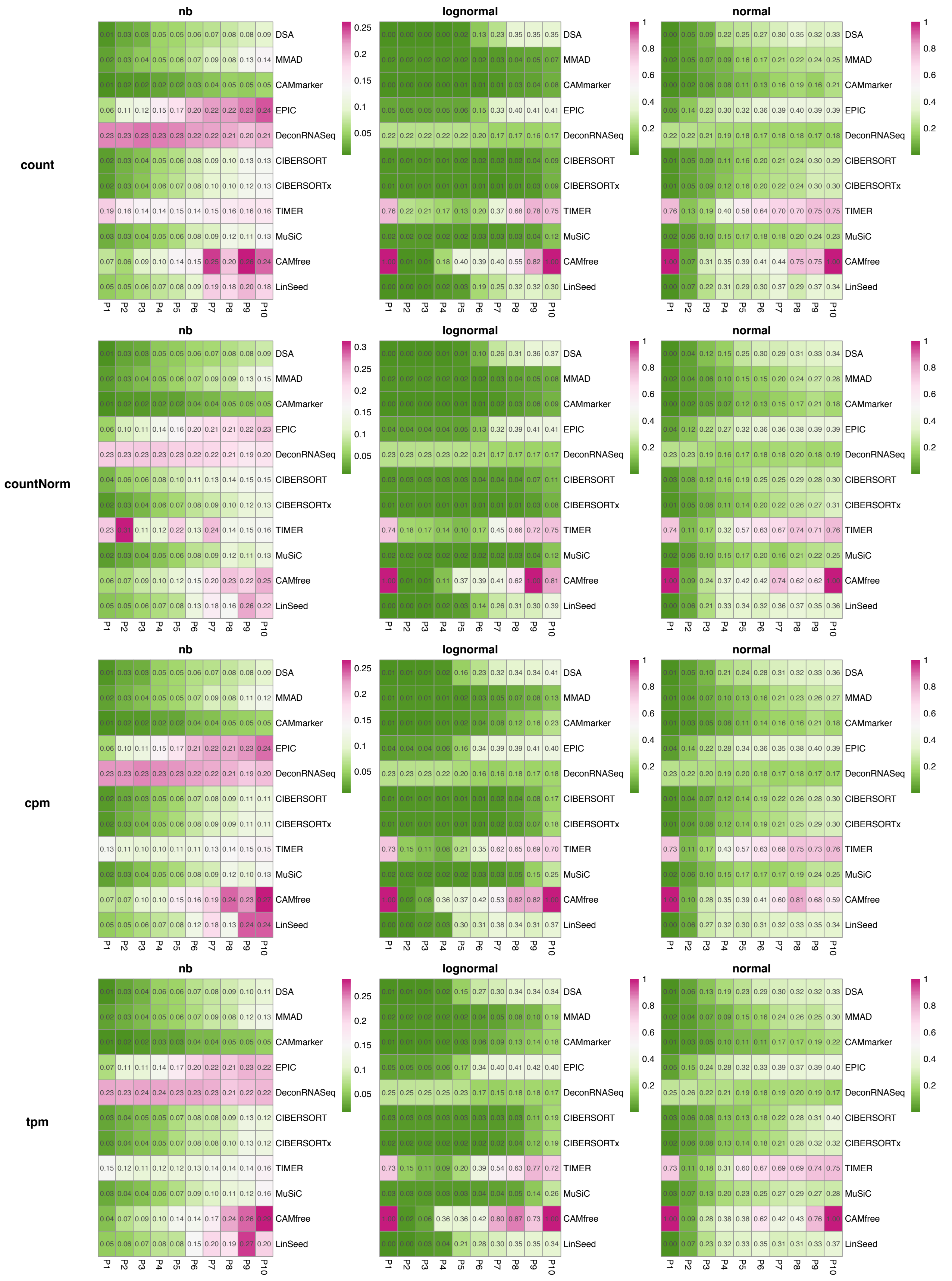
nb
lognormal
normal
count
countNorm
cpm
tpm
DSA
MMAD
CAMmarker
EPIC
DeconRNASeq
CIBERSORT
CIBERSORTx
TIMER
MuSiC
CAMfree
LinSeed
P1
P2
P3
P4
P5
P6
P7
P8
P9
P10

**Supplementary Fig. 4| mAD heatmap of sim1_simModel**

The deconvolution results of Sim1_simModel are organized by 12 heatmaps where each row panel refers to the results derived from the same quantification units and each column panel refers to the results derived from the same simulation model. In each heatmap, row indexes refer to the tested deconvolution methods and column indexes refer to the noise level. Each cell in the heatmap indicates averaged mADs of 9 testing sets (3 mixture sets × 3 reference sets)(Methods).

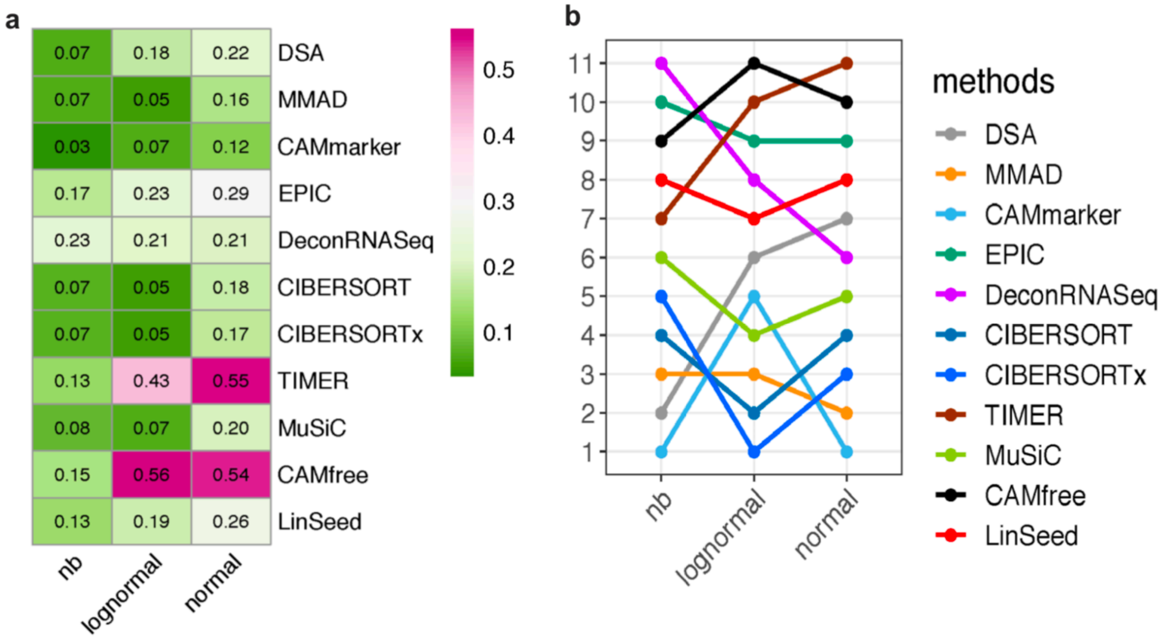


**Supplementary Fig. 5| Evaluation results of sim1_simModel based on mAD**

**a,** Heatmap of summarized evaluation results based on the mADs and **b,** rankings of tested deconvolution methods. In each heatmap, row indexes refer to the tested methods and column indexes refer to the simulation models (negative binomial, log-normal, and normal).

Variance Structure of GSE60424

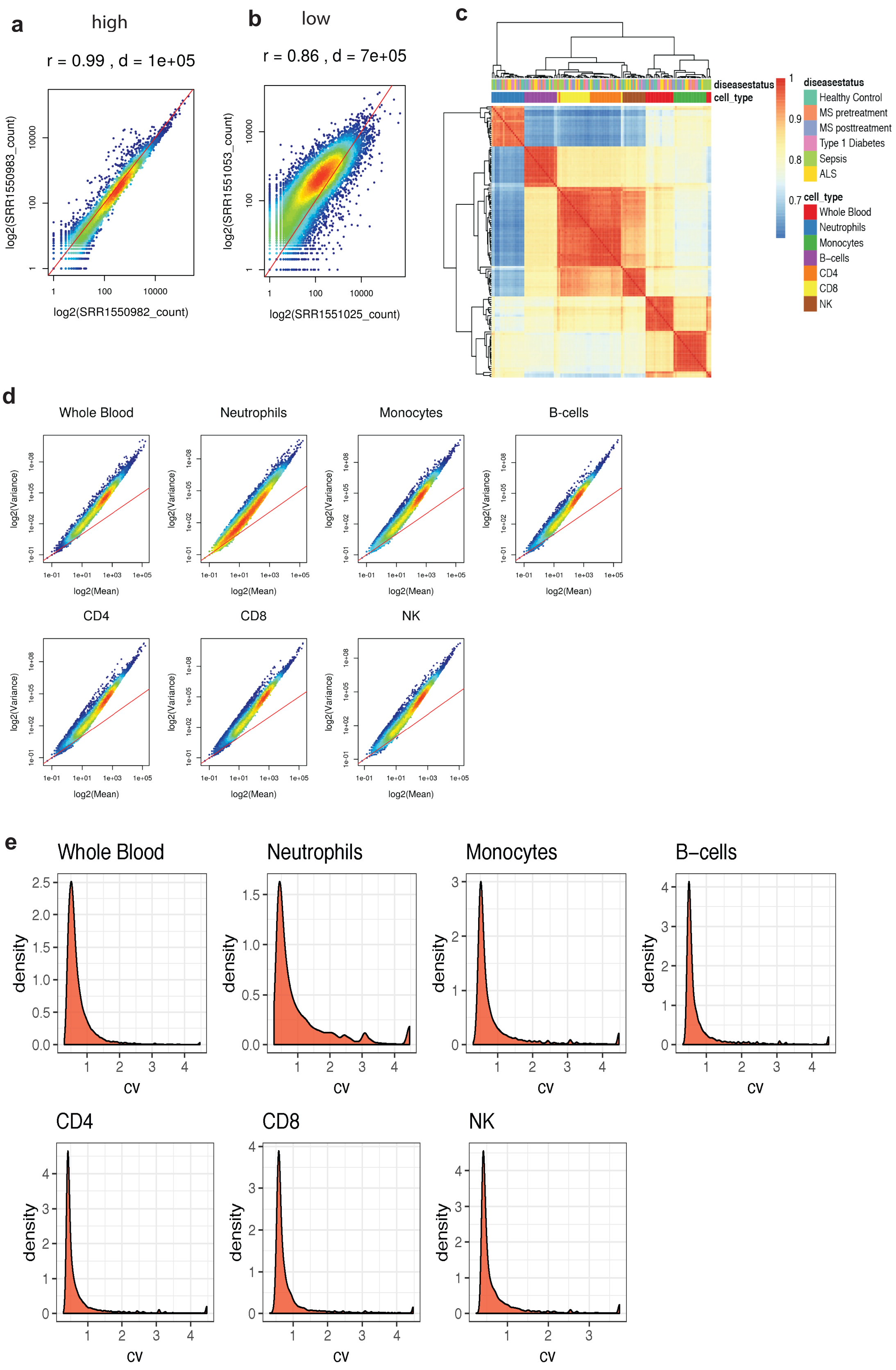

### Supplementary Fig. 6| Variance analysis of GSE60424

**a,b,** Scatter plots of whole blood sample profiles with (**a**) highest correlation and (**b**) lowest correlation. **c,** Correlation heatmap across all sample pairs. **d,** Mean-variance plot **e,** Density plot of CV (coefficient of variance) r: Spearman correlation coefficient, d: Euclidian distance, all points are plotted in the log space. (All results are in the count unit.)

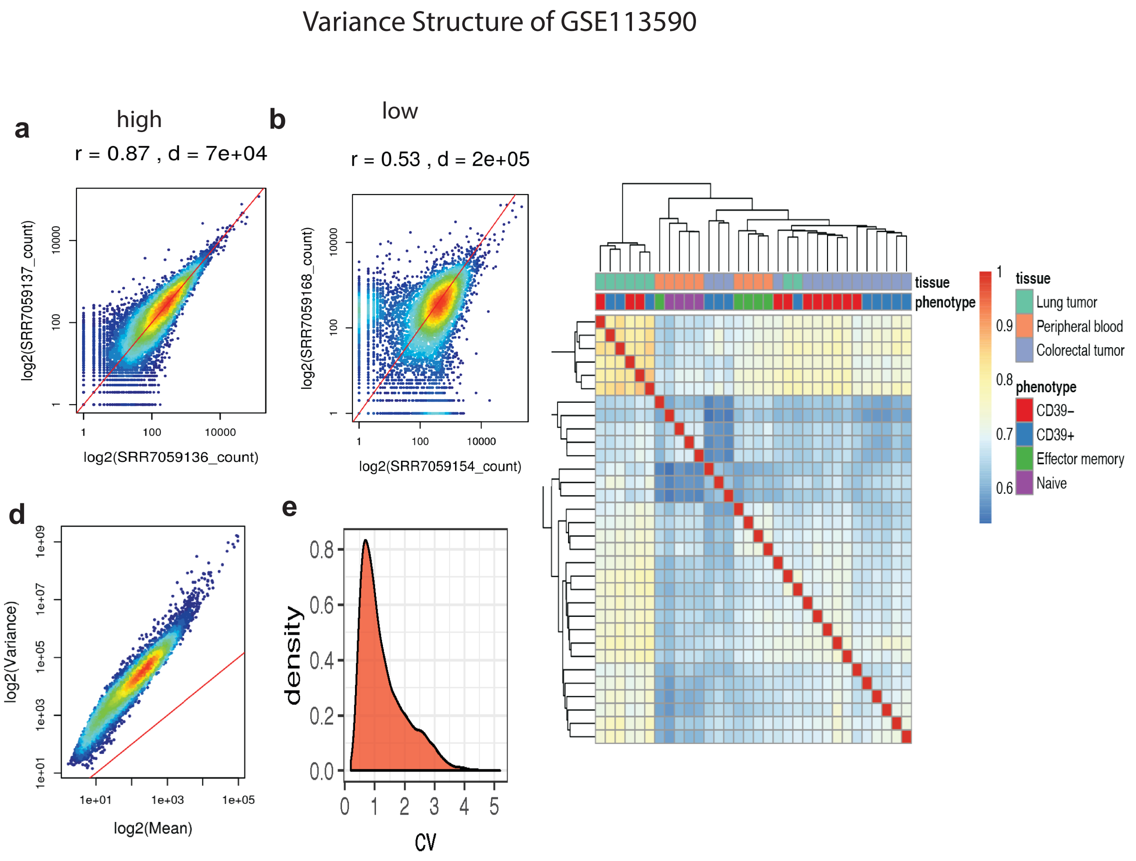


**Supplementary Fig. 7| Variance analysis of GSE113590**

**a,b,** Scatter plots of CD8 T sample profiles with (**a**) highest correlation and (**b**) lowest correlation. **c,** Correlation heatmap across all sample pairs. **d,** Mean-variance plot **e,** Density plot of CV (Coefficient of variance) r: Spearman correlation coefficient, d: Euclidian distance, all points are plotted in the log space. (All results are in the count unit.)

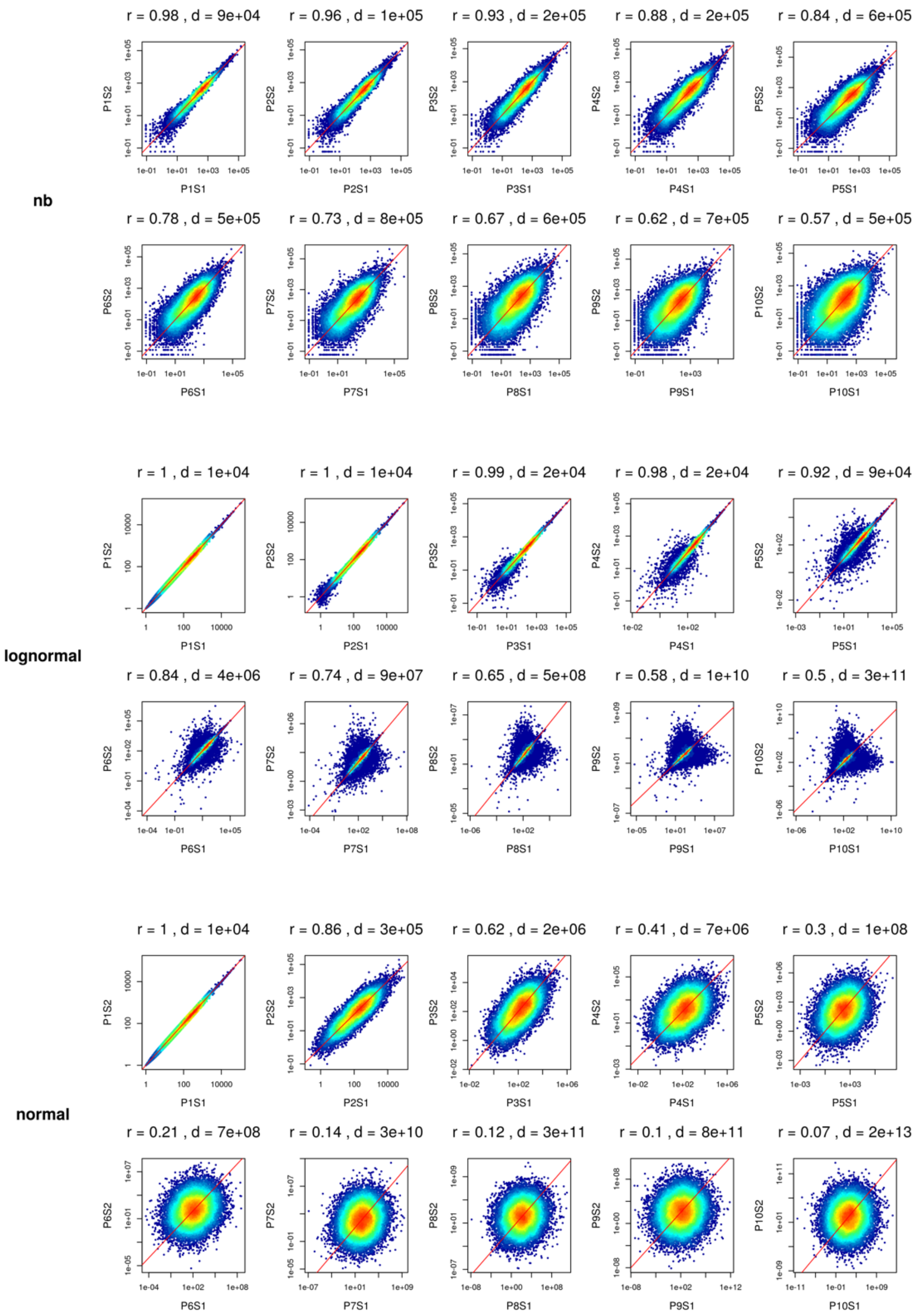


### Supplementary Fig. 8| Scatter plots of simulated profiles in Sim1_simModel

Sample-sample scatter plots of the first two simulated profiles that are derived from D1 (GSE51984). Each row panel refers to the simulation model used to generate simulation. (All results are in the count unit.)

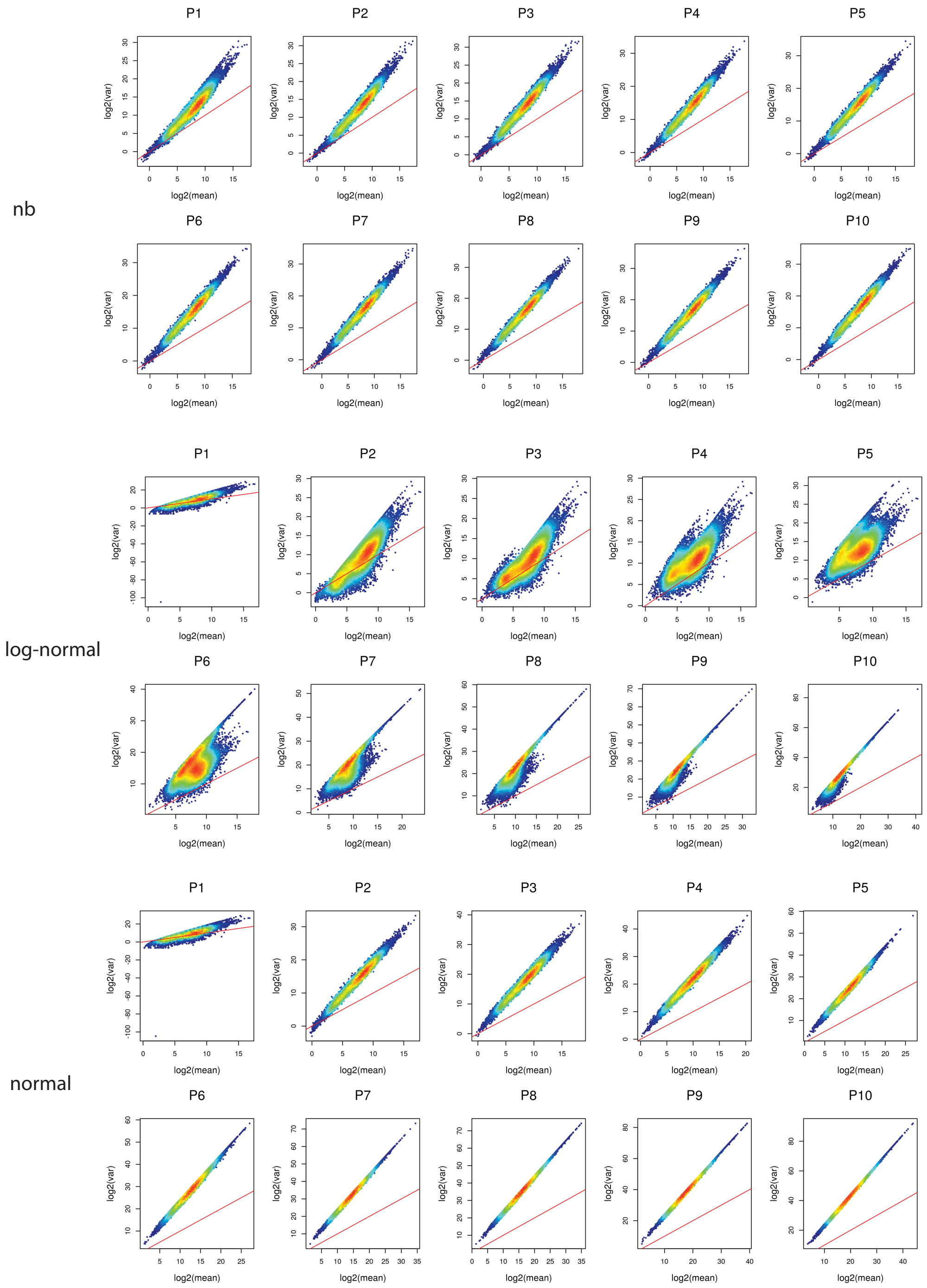


**Supplementary Fig. 9| Mean-variance plots of simulations in Sim1_simModel**

Mean-variance plots of simulations derived from negative binomial (nb), log-normal, and normal simulation models. In each simulation model, 10 plots are generated for 10 noise levels (P1–P10). The simulations in this figure are generated from GSE51984 (D1). (All results are in the count unit.)

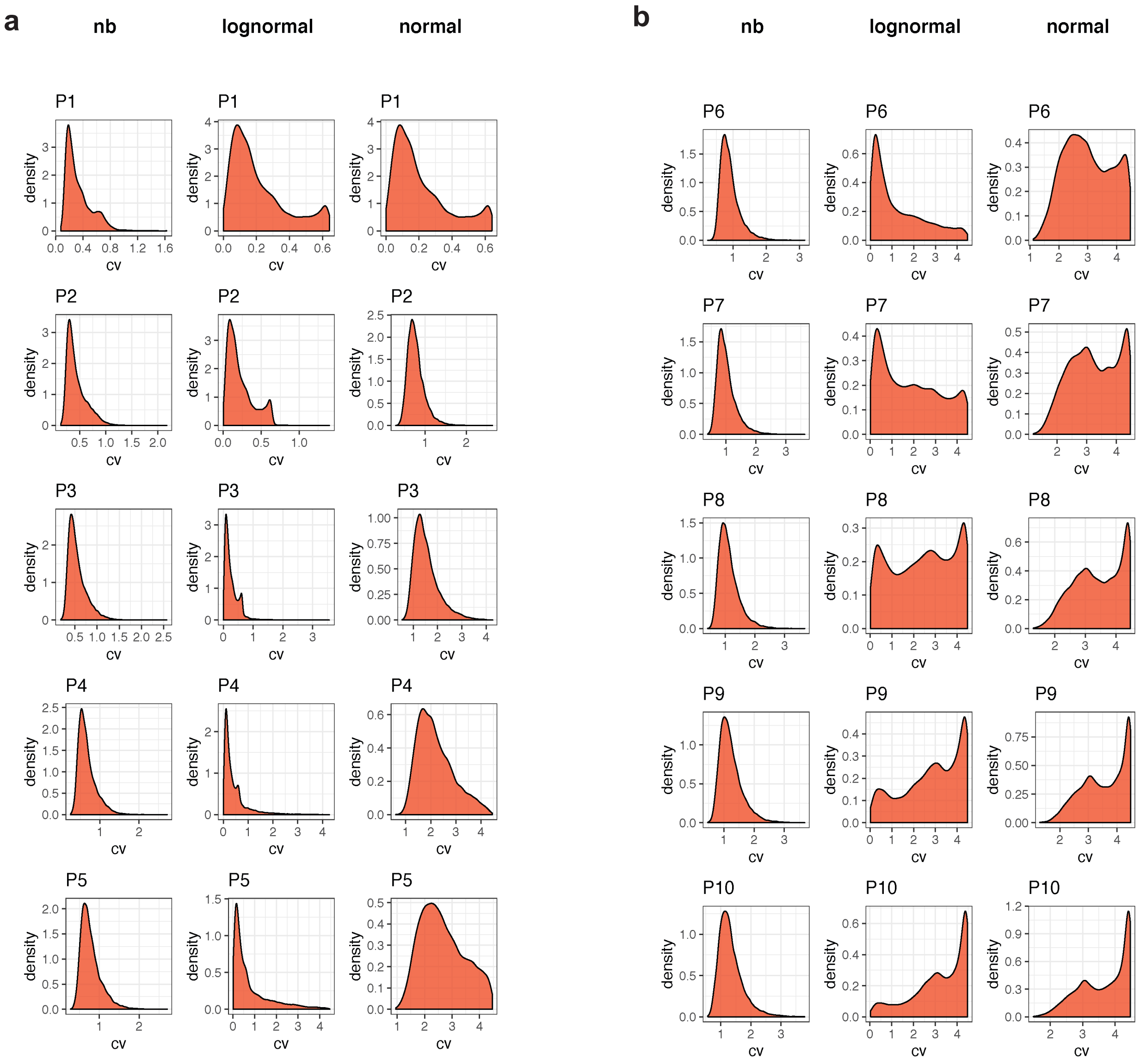


**Supplementary Fig. 10| Density plots of CV values in Sim1_simModel**

**a, b,** Density curve of CV (Coefficient of variance) from 3 simulation models with noise level (**a**) P1 – P5 and (**b**), P6 – P10. (All results are in the count unit.)

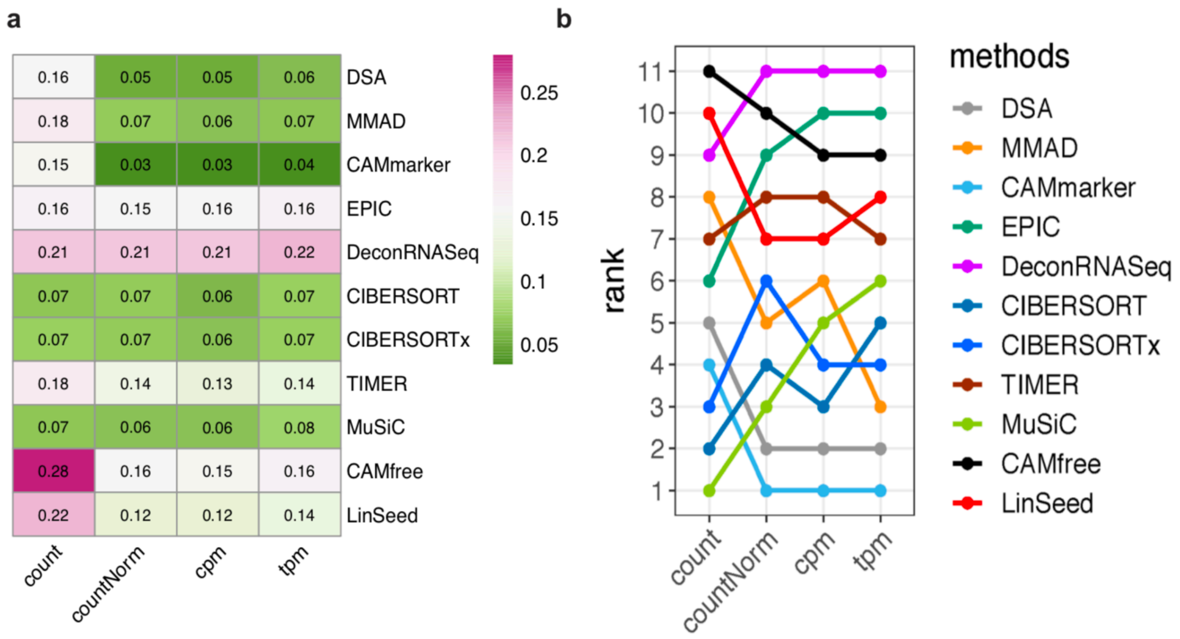


**Supplementary Fig. 11| Evaluation results of Sim1_libSzie**

**a,** Heatmap of summarized evaluation results based on the mADs and **b,** rankings of tested deconvolution methods. In each heatmap, row indexes refer to the tested methods and column indexes refer to the quantification units (count, countNorm, cpm, and tpm).

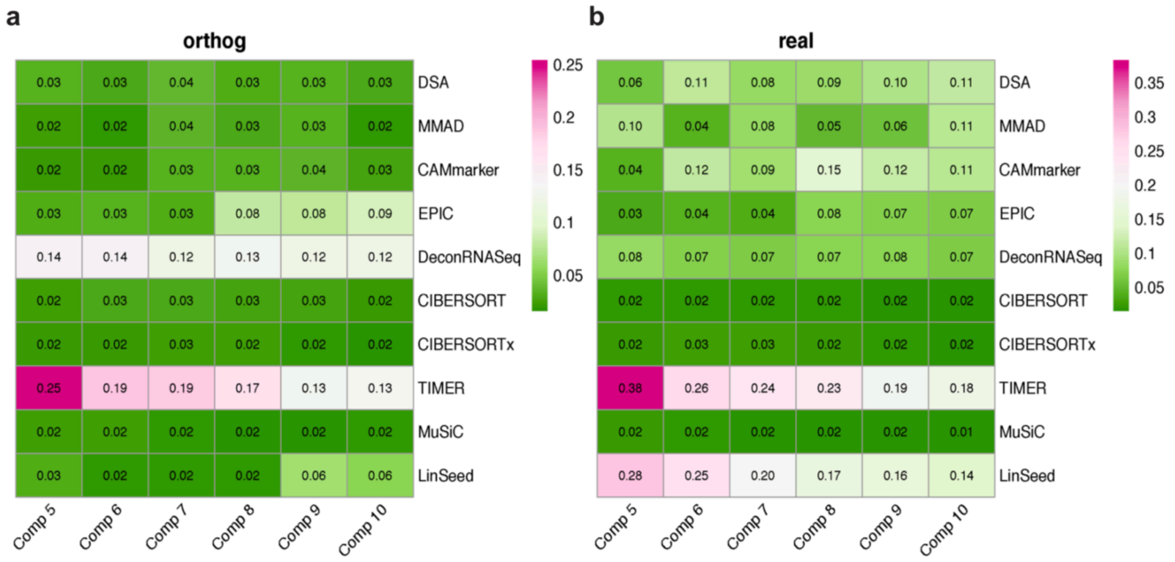


### Supplementary Fig. 12| Evaluation results of Sim2

**a,b,** Heatmaps of summarized evaluation results based on the mAD metric with (**a**) 'orthog' weight matrix and (**b**) 'real' weight matrix. In each heatmap, row indexes refer to the tested methods and column indexes refer to the cellular component numbers.

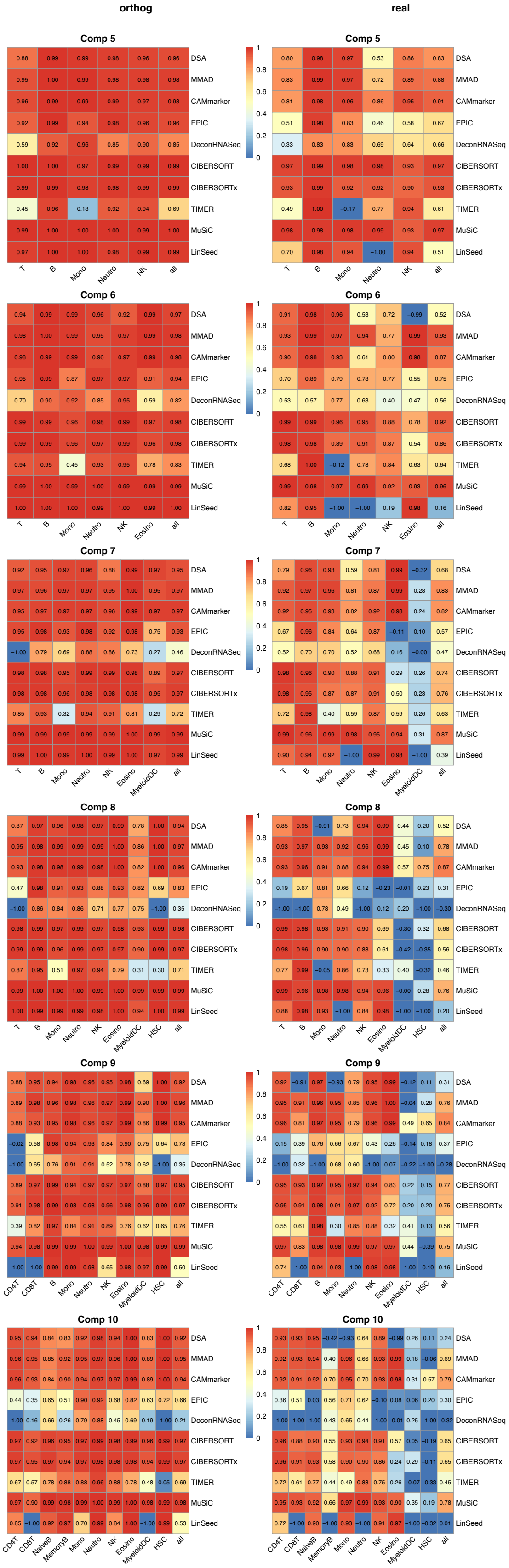
orthog
real
Comp 5
Comp 6
Comp 7
Comp 8
Comp 9
Comp 10
DSA
MMAD
CAMmarker
EPIC
DeconRNASeq
CIBERSORT
CIBERSORTx
TIMER
MuSiC
LinSeed

### Supplementary Fig. 13| Cell-type-specific correlation of Sim2

Evaluation results of Sim2 based on the correlation metric. Each row panel indicates the number of cellular components in the mixture and each column panel indicates the weight matrix type ('orthog' vs. 'real'). In each heatmap, row indexes refer to the tested methods, column indexes refer to the cell types, and the last column 'all' refers to the averaged evaluation results across all cell types.

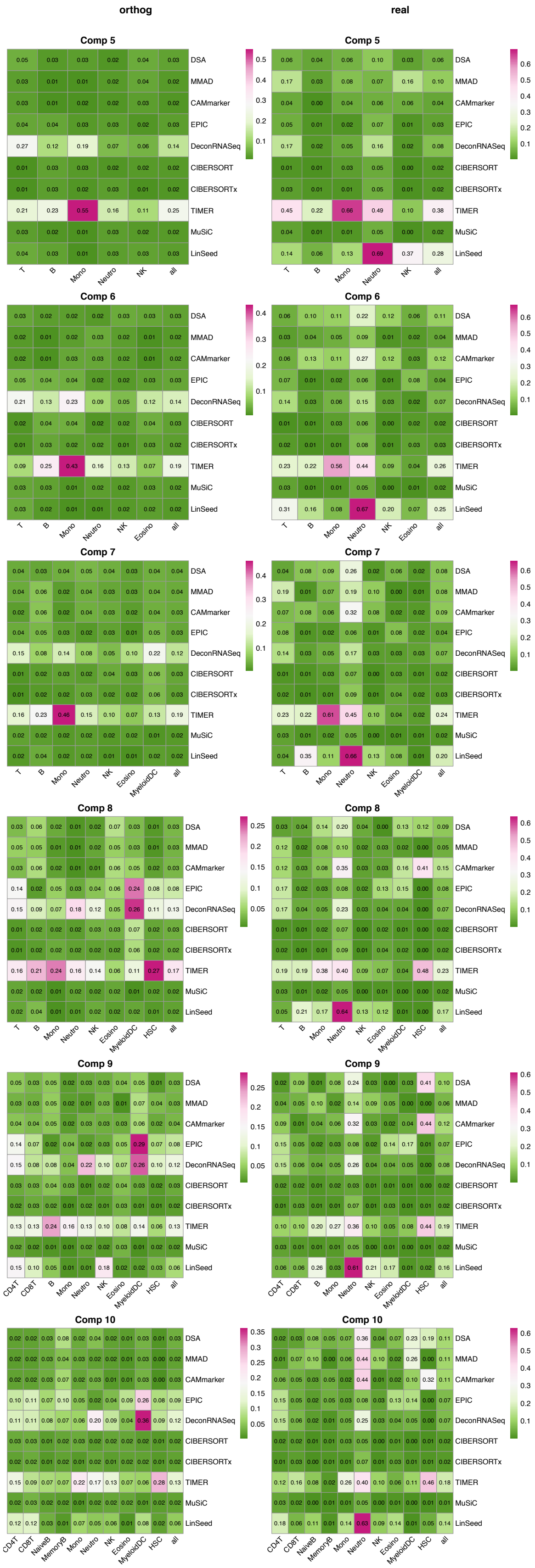
orthog
real
Comp 5
Comp 6
Comp 7
Comp 8
Comp 9
Comp 10
DSA
MMAD
CAMmarker
EPIC
DeconRNASeq
CIBERSORT
CIBERSORTx
TIMER
MuSiC
LinSeed
T
B
CD4T
CD8T
NaiveB
MemoryB
Mono
Neutro
NK
Eosino
MyeloidDC
HSC
all

### **Supplementary Fig. 14| Cell-type-specific mAD of Sim2**

Evaluation results of Sim2 based on the mAD metric. Each row panel indicates the number of cellular components in the mixture and each column panel indicates the weight matrix type (‘orthog’ vs. ‘real’). In each heatmap, row indexes refer to the tested methods, column indexes refer to the cell types, and the last column ‘all’ refers to the averaged evaluation results across all cell types.

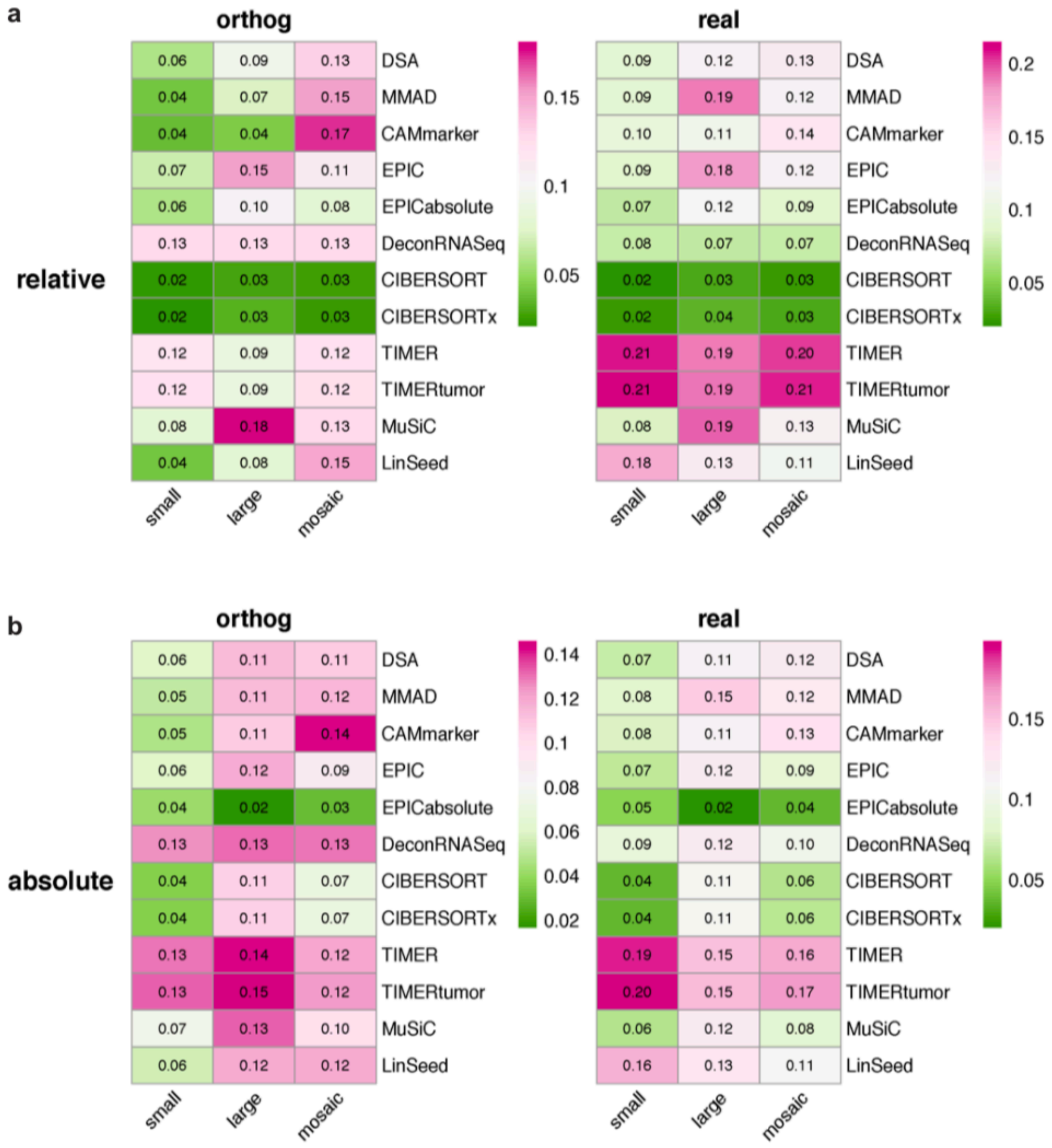


**Supplementary Fig. 15| Summarized mAD heatmaps of Sim3**

**a,b,** Heatmaps of summarized evaluation metric based on mAD evaluation metric with (**a**) relative measurement scale and (**b**) absolute measurement scale. In each heatmap, row indexes refer to the tested methods, and column indexes refer to the types of tumor spike-ins (small, large, and mosaic).

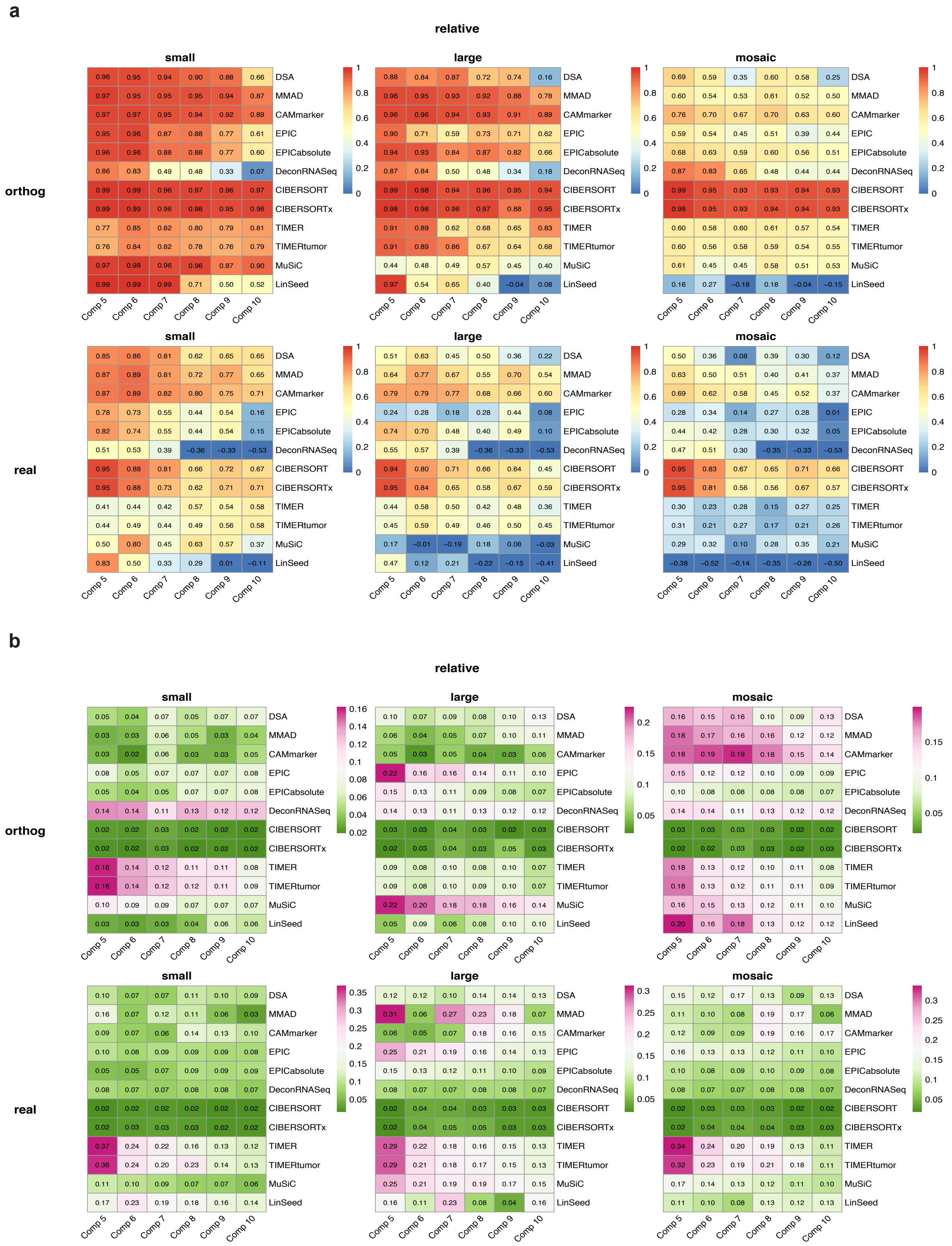
a
relative
small
large
mosaic
orthog
real
DSA
MMAD
CAMmarker
EPIC
EPICabsolute
DeconRNASeq
CIBERSORT
CIBERSORTx
TIMER
TIMERtumor
MuSiC
LinSeed
Comp 5
Comp 6
Comp 7
Comp 8
Comp 9
Comp 10
b
relative
small
large
mosaic
orthog
real

145

**Supplementary Fig. 16| Summarized evaluation heatmaps of Sim3 in the relative scale**

**a, b,** Summarized evaluation heatmap based on (**a**) Pearson's correlation coefficients and (**b**) mAD. The row panel indicates the type of weight matrix and the column panel indicates the type of tumor spike-ins (small, large, and mosaic). In each heatmap, row indexes refer to the tested methods and column indexes refer to the number of cellular components in the mixture.

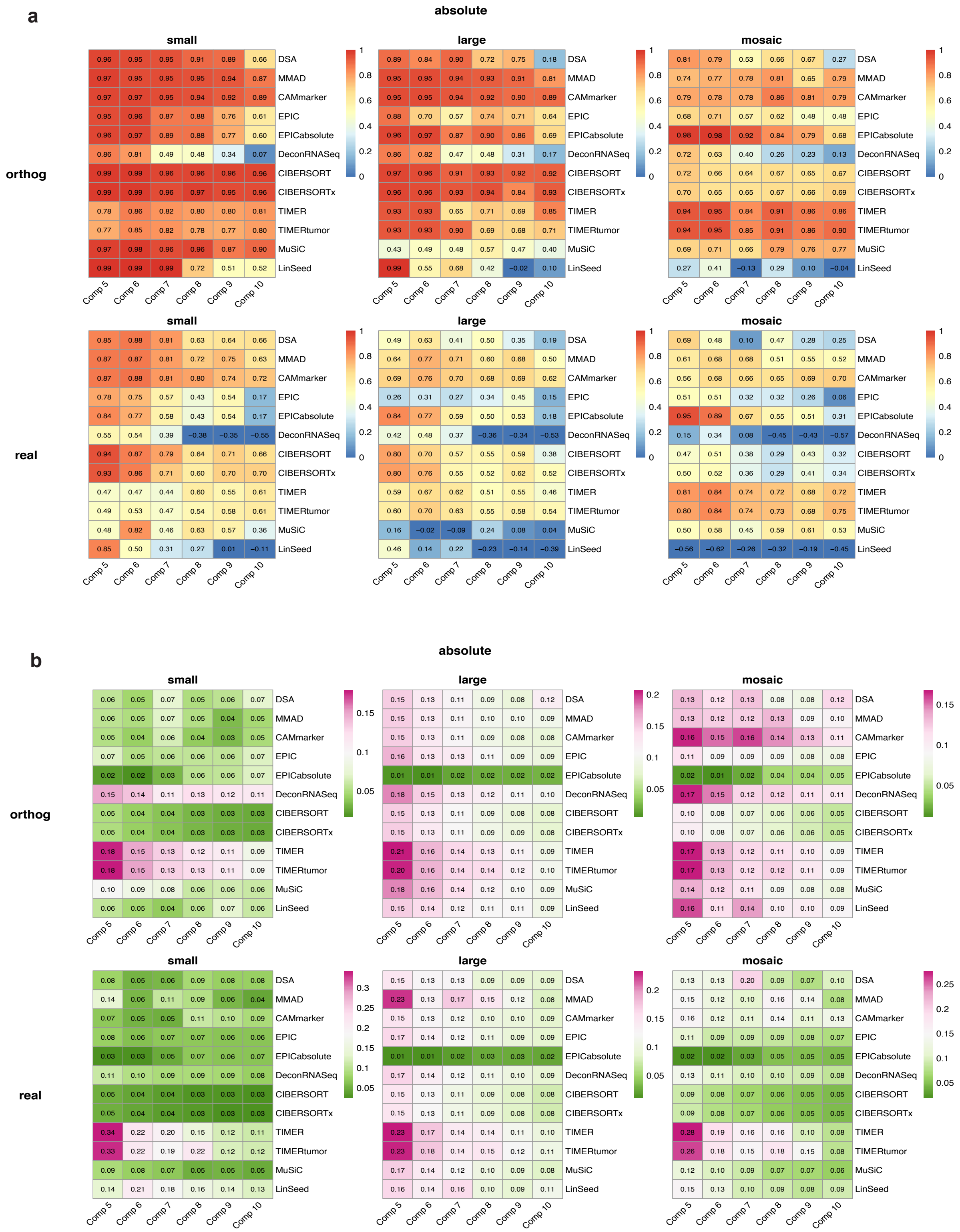
a
absolute
small
large
mosaic
orthog
real
b
DSA
MMAD
CAMmarker
EPIC
EPICabsolute
DeconRNASeq
CIBERSORT
CIBERSORTx
TIMER
TIMERtumor
MuSiC
LinSeed
Comp 5
Comp 6
Comp 7
Comp 8
Comp 9
Comp 10

**Supplementary Fig. 17| Summarized evaluation heatmaps of Sim3 in the absolute scale**

**a, b,** Summarized evaluation heatmap based on (**a**) Pearson's correlation coefficients and (**b**) mAD. The row panel indicates the type of weight matrix and the column panel indicates tumor spike-ins (small, large, and mosaic). In each heatmap, row indexes refer to the tested methods and column indexes refer to the number of cellular components in the mixture.
